# Non cell-autonomous effect of astrocytes on cerebral cavernous malformations

**DOI:** 10.1101/2021.01.29.428891

**Authors:** Miguel Alejandro Lopez-Ramirez, Shady Ibrahim Soliman, Preston Hale, Catherine Chinhchu Lai, Angela Pham, Esau Estrada, Sara McCurdy, Romuald Girard, Riya Verma, Thomas Moore, Rhonda Lightle, Nicholas Hobson, Robert Shenkar, Orit Poulsen, Gabriel G. Haddad, Richard Daneman, Brendan Gongol, Hao Sun, Frederic Lagarrigue, Issam A. Awad, Mark H. Ginsberg

## Abstract

Cerebral cavernous malformations (CCMs) are common neurovascular lesions caused by loss-of-function mutations in one of three genes, including *KRIT1* (CCM1), *CCM2*, and *PDCD10* (CCM3), and generally regarded as an endothelial cell-autonomous disease. Here we report that proliferative astrocytes play a critical role in CCM pathogenesis by serving as a major source of VEGF during CCM lesion formation. An increase in astrocyte VEGF synthesis is driven by endothelial nitric oxide (NO) generated as a consequence of KLF2 and KLF4-dependent elevation of eNOS in CCM endothelium. The increased brain endothelial production of NO stabilizes HIF-1α in astrocytes, resulting in increased VEGF production and expression of a “hypoxic” program under normoxic conditions. We show that the upregulation of cyclooxygenase-2 (COX-2), a direct HIF-1α target gene and a known component of the hypoxic program, contributes to the development of CCM lesions because the administration of a COX-2 inhibitor significantly prevents progression of CCM lesions. Thus, non-cell-autonomous crosstalk between CCM endothelium and astrocytes propels vascular lesion development, and components of the hypoxic program represent potential therapeutic targets for CCMs.

## Introduction

Cerebral cavernous malformations (CCMs) are caused by gross brain endothelial changes that lead to blood-brain barrier (BBB) dysfunction, resulting in significant morbidity and mortality(1, 2). CCMs affect ~1/200 children and adults, causing a lifelong risk of chronic and acute hemorrhage and consequent disabilities from neurological deficits, for which there is no current effective pharmacologic therapy(3–6). Inherited germline and somatic loss of function mutations in the genes *KRIT1* (Krev1 interaction trapped gene 1, *CCM1), CCM2 (Malcavernin), PDCD10* (Programmed cell death protein 10, *CCM3*) are associated with CCMs (5, 7–9). Notably, CCMs have been recognized as an endothelial cell-autonomous disease because endothelial-specific inactivation of murine *Krit1, Ccm2 or Pdcd10*, results in brain and retinal vascular lesions similar to those in CCM patients(10–13). These murine studies are also supported by the finding of a “second hit” on the normal *KRIT1* or *PDCD10* allele in the CCM endothelium (14, 15). However, there has been to date no compelling mechanistic explanation for the propensity of CCM lesions to form in the central nervous system (CNS).

Recent studies have shown that loss of *CCM* genes in neuroglia or stromal cells may contribute to CCM pathogenesis through non-cell-autonomous mechanisms (16–18). Although CCM is a disease that affects the neurovascular unit, there is a limited understanding of the crosstalk between the CCM endothelium and astrocytes, (17, 19, 20). Astrocytes are the most abundant cell in the CNS and form part of the neurovascular unit that maintains a functional BBB(21–23). Additionally, astrocytes respond to multiple forms of insults and diseases by a process called reactive astrogliosis or astrocytosis, which involves changes in morphology, function and gene expression (24, 25). Reactive astrogliosis increases secretion of VEGF and can contribute to the development and progression of CNS disease (26–28). Previous studies have demonstrated elevated levels of angiogenic factor VEGF in CCM lesions, and in the plasma of individuals with the hereditary and sporadic form of the disease(29–32). VEGF signaling mediates disruption of the brain endothelial barrier by disassembly of inter-endothelial junctions(33) and can cause hemorrhages(34, 35), both prominent features of CCM lesions(2, 6, 10, 36, 37). We and others found that CCM led to a dramatic increase in VEGF signaling and that blocking VEGF signaling prevents subsequent formation of CCMs(6, 38). However, despite these emerging studies, the mechanism and source of VEGF production in CCM lesions in vivo remain elusive.

Endothelial nitric oxide synthase (eNOS, the product of the *NOS3* gene) is an enzyme that generates nitric oxide (NO) and it is responsible for vascular remodeling, angiogenesis and vascular tone(39, 40). Previous studies indicate that eNOS contributes to VEGF-induced angiogenesis and vascular permeability(41) through NO production (42, 43). Here we show that astrocytes are a major source of VEGF during CCM pathogenesis and that depletion of proliferative astrocytes prevents CCM lesion development. We show that CCM endothelium-induced elevation of astrocyte-derived VEGF in neonatal and juvenile brains occurs through elevation of brain eNOS/NO-dependent signaling. Production of NO in CCM endothelium contributes to normoxic stability of HIF-1α in astrocytes. Our findings indicate that the eNOS increase in human CCM endothelium is ascribable to the elevation of the transcription factors KLF2 and KLF4, previously implicated in the CCM pathogenesis(6, 13, 37, 44, 45). Furthermore, genetic inactivation of one copy of the *Nos3* gene is sufficient to attenuate CCM endothelial NO production, HIF-1α protein stability and astrocyte-derived VEGF expression. Our results further reveal that CCM tissue results in increased HIF activity under normoxic conditions. We show that pharmacological inhibition of a HIF-1α regulated gene, cyclooxygenase-2, ameliorates the development of CCM lesions in animal models. Overall, these findings indicate that a non-cell-autonomous mechanism mediated by CCM endothelium-driven normoxic stabilization of HIF-1α and increase of VEGF in astrocytes makes a major contribution to CCM pathogenesis. Understanding the crosstalk between dysfunctional brain vasculature and components of the neurovascular unit (e.g., astrocytes) has the potential to lead to the development of novel therapeutic strategies for CCM disease as exemplified by our finding that inhibition of HIF1-driven COX-2 by an approved and a well-tolerated drug can ameliorate murine models of CCM disease.

## Results

### Astrocytes contribute to cerebral cavernous malformations development

CCMs have been studied as an endothelial cell-autonomous disease (6, 14, 15), marked by changes in the CNS vasculature due to the loss of endothelial *PDCD10, KRIT1* or *CCM2* (10–12). Although it is a disease that affects the neurovascular unit, we know little about whether astrocytes influence CCM pathogenesis(17, 19). This prompted us to investigate the relationship between astrocytes and CCM lesion development in murine CCMs (Fig. 1A,B, fig. S1). Neonatal endothelial-specific inactivation of murine *Pdcd10*, producing *Pdcd10^ECKO^* mice(37), results in cerebellar vascular lesions detected in sections stained by hematoxylin and eosin (Fig. 1A,B, fig. S1) or by observing dramatic vascular dilation by staining the endothelial marker CD34 at P9 (Fig. 1A,B). CCM lesions have high propensity to develop in GFAP-positive fibrous astrocytes(46) (Fig. 1A) in the white matter of *Pdcd10^ECKO^* hindbrains (fig. S1). Moreover, P8 *Krit1^ECKO^* hindbrains show the same spatial distribution of CCM lesions among astrocytes positive for GFAP on the white matter tract (fig. S1). To examine the role of astrocytes, we generated *Pdcd10^ECKO^;GFAP-TK* mice *(Pdgfb-iCreERT2;Pdcd10^fl/fl^; GFAP-TK (Thymidine kinase)* mice) in which proliferative astrocytes are selectively depleted in a time-controlled manner, by administration of antiviral agent ganciclovir (GCV)(47, 48). Administration of 20 mg/Kg of GCV to neonatal mice at P7 markedly reduced hindbrain vascular lesions in P9 *Pdcd10^ECKO^;GFAP-TK* mice when compared with GCV-treated littermate *Pdcd10^ECKO^* controls (Fig. 1C). To quantify CCM formation, we imaged P9 hindbrains using contrast-enhanced, high-resolution x-ray micro-computed tomography (micro-CT), and measured lesion volumes using semiautomated software(49). Blinded measurement of total CCM lesion volume showed that GCV-treated *Pdcd10^ECKO^;GFAP-TK* mouse hindbrains exhibited significant reduction of CCM lesions compared with GCV-treated *Pdcd10^ECKO^* mouse hindbrains littermates (Fig. 1D). These data suggest that in CCM disease, proliferative astrocytes participate in vascular lesions formation.

**Fig 1.**
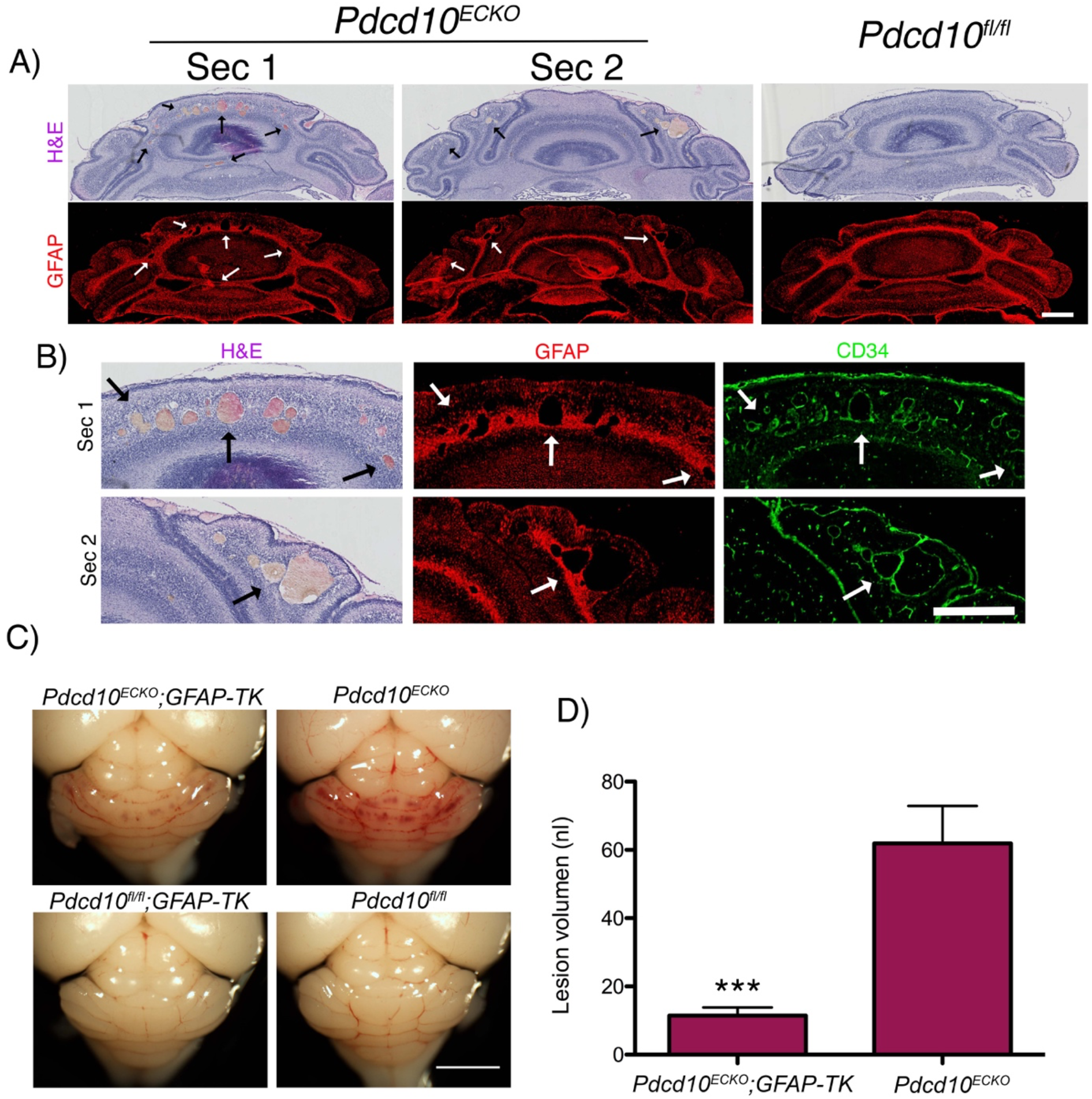
Astrocytes contribute to cerebral cavernous malformations development. (**A**)Histological analysis of cerebellar sections from P9 *Pdcd10^ECKO^* and littermate control *Pdcd10^fl/fl^* mice. Low magnification of CCM lesions detected in sections stained by hematoxylin and eosin. CCM lesions spatially developed on fibrous astrocyte areas positive for GFAP immunostaining (red). Arrows indicate CCM lesions. (**B**) Magnified region from section 1 and 2 (Sec1 and Sec2) from *Pdcd10^ECKO^mice* in *A*, dramatic vascular dilation is shown with immunohistochemistry for the endothelial marker CD34 (green). Arrows indicate CCM lesions. (**C**) Administration of 1.5 mg/Kg of GCV in neonatal at P7 markedly reduced vascular lesions in P9 *Pdcd10^ECKO^;GFAP-TK* hindbrains when compared with GCV-treated littermate *Pdcd10^ECKO^* control. (**D**) Quantification of lesion volumes by micro-CT analysis from mice in experiments showed in *C*. *n*= 8 or 9 mice in each group. Data are mean±SEM. ***, P<0.001; determine by Student’s *t* test. Scale bars: (**A** and **B**) 500 μm; (**C**) 2 mm.

### Astrocyte-derived VEGF is increased during CCM development

CCMs exhibit increased VEGF signaling(6, 30, 31) due to the loss of an anti-angiogenic checkpoint(6). To ascertain a potential source of increased expression of VEGF, we introduced a VEGF reporter, *Vegfa^tm1.1Nagy^*, in CCM animal models. *Vegfa^tm1.1Nagy^* carries a nuclear-localized beta-galactosidase (β-gal) knock-in at the 3′ UTR of the *Vegfa* gene locus that permits singlecell monitoring of VEGF expression(50). We observed an increase in β-gal/VEGF expression, as shown by X-gal staining, in areas surrounding CCM lesions in P10 *Pdcd10^ECKO^; Vegfa^tm1.1Nagy^* hindbrains when compared with *Pdcd10^fl/fl^; Vegfa^tm1.1Nagy^* littermate control (Fig. 2A). We found that most β-gal/VEGF+ cells colocalized with the GFAP marker of fibrous astrocytes and Bergman glia located within Purkinje cell layer(46) in *Pdcd10^ECKO^; Vegfa^tm1.1Nagy^* (Fig. 2A). We also observed that calbidin-positive Purkinje neurons exhibited increased β-gal/VEGF expression near CCM lesions (Data not shown). Consistent with results observed using β-gal/VEGF reporter mice, high-resolution in situ hybridization (ISH) analysis confirmed an increase in VEGF expression near and around the CCM lesion colocalized with the GFAP marker of astrocytes (Fig. 2B, fig. S2). Moreover, we observed that *Pdcd10^ECKO^* hindbrains exhibited ~1.8-fold increase in *Vegfa* mRNA levels compared with littermate *Pdcd10^fl/fl^*control (Fig. 2C). Administration of GCV to neonatal mice at P7 significantly reduced *Vegfa* mRNA levels in vascular lesions in P10 *Pdcd10^ECKO^;GFAP-TK* mice compared with GCV-treated littermate *Pdcd10^ECKO^* controls (Fig. 2C). We next tested the effect of depletion of proliferative astrocytes at the developmental stage when vascular lesions are present in *Pdcd10^ECKO^*. Our data indicate that administration of GCV to neonatal mice between P9 to P12 significantly reduced *Vegfa* mRNA levels and CCM lesion burden in P15 *Pdcd10^ECKO^;GFAP-TK* mice when compared with GCV-treated littermate *Pdcd10^ECKO^* controls (Fig. 2D,E). Since the retinal blood vessels is another vascular bed affected by the loss of *Pdcd10* in endothelial cells, we also compared the subcellular expression of VEGF. Deletion of the endothelial *Pdcd10* gene at P3 prominently increase β-gal/VEGF expression in GFAP+ retinal astrocytes at the leading edge of blood vessel growth and immediately ahead of the plexus in P9 *Pdcd10^ECKO^; Vegfa^tm1.1Nagy^* retinas (fig. S2,S3). We observed that the enhanced expression of β-gal/VEGF was associated with impaired extension of the developing superficial vascular plexus(51) in P9 *Pdcd10^ECKO^; Vegfa^tm1.1Nagy^* retinas when compared with *Pdcd10^fl/fl^; Vegfa^tm1.1Nagy^* littermate control (fig. S3). Moreover, defect on the extension of the vascular plexus in P12 *Pdcd10^ECKO^; Vegfa^tm1.1Nagy^* retinas was reduced compared with *Pdcd10^fl/fl^; Vegfa^tm1.1Nagy^* littermate control, whereas the endothelial density and an increase in β-gal/VEGF expression in GFAP+ retinal astrocytes were elevated (fig. S3). These data suggest that increased expression of astrocyte-derived VEGF could account for a contribution of astrocytes to CCM lesion formation.

**Fig 2.**
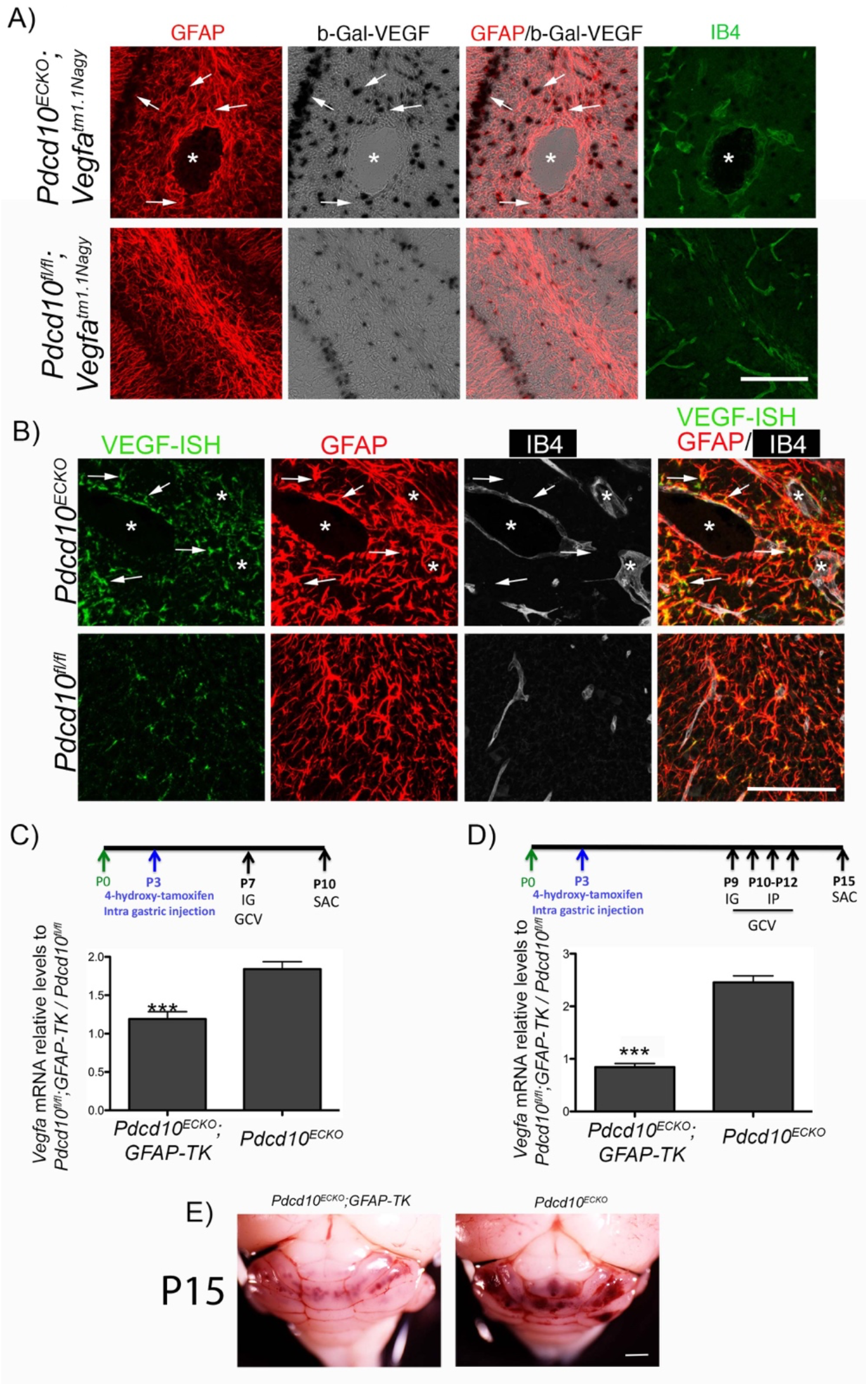
Astrocyte-derived VEGF increased during cerebral cavernous malformations development. (**A**) Confocal microscopy of cerebellar cortex from P10 *Pdcd10^ECKO^* and littermate control *Pdcd10^fl/fl^* mice stained for GFAP-positive astrocytes (red), β-gal/VEGF expression detected by X-gal staining (black), endothelial marker isolectin B4 (green). Asterisk, vascular lumen of CCM lesion. Arrows indicate β-gal/VEGF in GFAP-positive astrocytes (n=4). (**B**) ISH for VEGF (green) combined with immunohistochemistry to identify GFAP-positive astrocytes (red), endothelial marker isolectin B4 (IB4; white). Asterisk, vascular lumen of CCM lesion. Arrows indicate VEGF and GFAP-positive astrocytes co-localization (n=2). (**C**) Quantification of *Vegfa* mRNA levels n P10 *Pdcd10^ECKO^;GFAP-TK* and *Pdcd10^ECKO^* hindbrains when compared with littermate *Pdcd10^fl/fl^;GFAP-TK or Pdcd10^fl/fl^* control respectively, as assessed by RT-qPCR. All mice received IG administration of 1.5 mg/Kg of GCV in neonatal at P7 (SEM, *n*=6 mice in each group). (**D**) Quantification of *Vegfa* mRNA levels in P15 *Pdcd10^ECKO^;GFAP-TK* and *Pdcd10^ECKO^* hindbrains when compared with littermate *Pdcd10^fl/fl^;GFAP-TK or Pdcd10^fl/fl^* control respectively, as assessed by RT-qPCR. All mice received IG administration of 1.5 mg/Kg of GCV in neonatal at P9 and three consecutive doses (P10 to P12) of IP 1.5 mg/Kg of GCV (SEM, *n*=5 or 6 mice in each group). (**E**) Prominent lesions are present in the cerebellum of P15 GCV-treated *Pdcd10^ECKO^* mice, whereas extensive reduction in lesions are observed in GCV-treated *Pdcd10^ECKO^;GFAP-TK* mice littermate (n=6 or 7). Data are mean±SEM. ***, P<0.001 (comparison to *Pdcd10^ECKO^* littermates); determine by Student’s *t* test. Scale bars: (**A**, and **B**) 100 μm.

### Astrocyte-derived VEGF increases during CCM formation in juvenile mice

The previous studies showed increased VEGF expression in an acute neonatal CCM model. Recent studies have emphasized that chronic models may be more reflective of the course of the human disease(52–54). We therefore, investigated the expression of VEGF increased in perilesional astrocytes in juvenile *Pdcd10^ECKO^;Vegfa^tm1.1Nagy^* mice. Similar to neonatal mice, we observed an increase in β-gal/VEGF expression in areas surrounding CCM lesions in P30 *Pdcd10^ECKO^; Vegfa^tm1.1Nagy^* cerebellar tissue (Fig. 3A). We found that most β-gal/VEGF+ cells colocalized with the GFAP astrocyte marker in *Pdcd10^ECKO^; Vegfa^tm1.1Nagy^* mice (Fig. 3A). Consistent with results observed using β-gal/VEGF reporter mice, *Pdcd10^ECKO^* cerebellar tissue showed a significant increase in *Vegfa* mRNA levels when compared with littermate *Pdcd10^fl/fl^* control (Fig. 3B). In contrast to neonatal *Pdcd10^ECKO^* mice, *Pdcd10^ECKO^* juvenile mice develop CCM lesions in the cerebrum(52, 53, 55) and these were accompanied by a marked increase in β-gal/VEGF expression in GFAP+ astrocytes and in SOX-9+ astrocytes in P30 *Pdcd10^ECKO^; Vegfa^tm1.1Nagy^* brains (Fig. 3A, fig. S3). We also observed a significant increase in *Vegfa* mRNA in the cerebrum of *Pdcd10^ECKO^* mice (Fig. 3C). Additionally, we found elevated levels of VEGF in plasma from P30 *Pdcd10^ECKO^* mice when compared with *Pdcd10^fl/fl^* littermate control (Fig. 3D). These data support the idea that astrocyte-derived VEGF contributes to CCM lesions(29–32).

**Fig 3.**
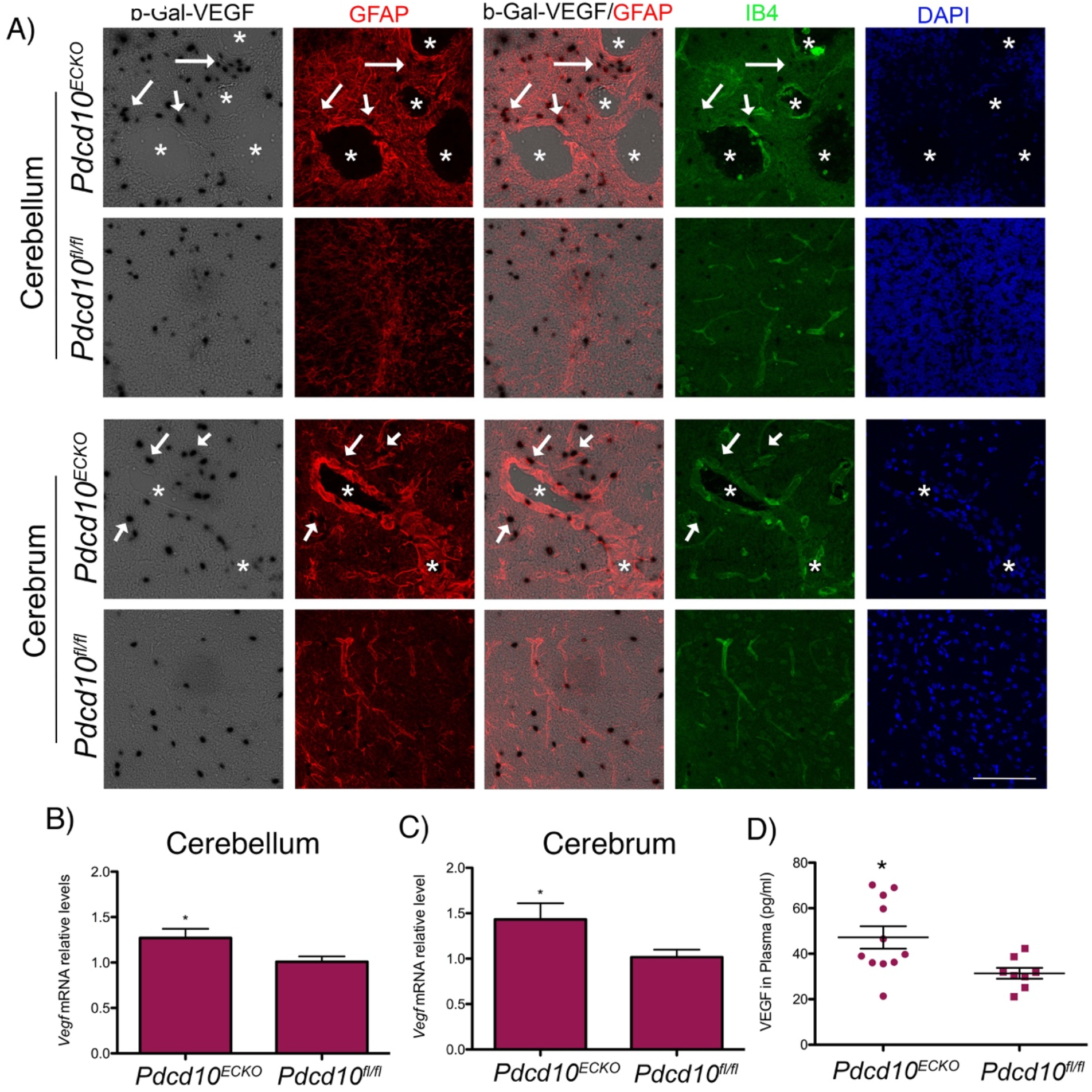
Astrocyte-derived VEGF increased during cerebral cavernous malformations in juvenile animal brain. (**A**) Confocal microscopy of brain (cerebellum and cerebrum cortex) from *Pdcd10^ECKO^;Vegfa^tm1.1Nagy^* and *Pdcd10^fl/fl^;Vegfa^tm1.1Nagy^* littermate control stained for β-gal/VEGF expression detected by X-gal staining (black), GFAP-positive astrocytes (red), isolectin B4 (green), and DAPI for nuclear DNA (blue). Asterisks, vascular lumen of CCM lesions. Arrows indicate β-gal/VEGF in GFAP-positive astrocytes. (**B**-**C**) Quantification of *Vegfa* mRNA levels in *Pdcd10^ECKO^* brains when compared with littermate *Pdcd10^fl/fl^*control, as assessed by RT-qPCR (SEM, *n*=4 or 6 mice in each group). (**D**) VEGF levels in plasma from *Pdcd10^ECKO^* mice and littermate *Pdcd10^fl/fl^* controls, as assessed by ELISA. *n*= 8 or 11 mice in each group. Data are mean±SEM. *, P<0.05; determine by Student’s *t* test. Scale bar: (**A**) 100 μm.

### Increase in normoxic HIF-1α stabilization in CCM astrocytes

Transcription factor HIF-1α regulates expression of VEGF(27, 56–58). Therefore, we next investigated whether the significant increase of VEGF in astrocytes during CCMs is due to elevation of HIF-1α. For this studies, we first prepared co-culture studies to better understand the interaction between CCM endothelium and astrocytes (Fig. 4A) and the level of HIF-1α in purified astrocytes (fig.S4) co-cultured with *Pdcd10^ECKO^* or *Pdcd10^fl/fl^* BMECs was determined. Immunocytochemistry revealed that the level of HIF-1α was elevated in astrocytes co-cultured with *Pdcd10^ECKO^* BMECs compared to those co-cultured with control *Pdcd10^fl/fl^* BMECs (Fig. 4A,B, fig. S4). Consistent with this observation, Western blot analysis of *Pdcd10^ECKO^* hindbrains at P10 confirmed an increase in HIF-1α protein levels when compared with *Pdcd10^fl/fl^* litermate controls (Fig. 4C). Furthermore, we observed an increase in HIF-1α activity in *Pdcd10^ECKO^* hindbrains (Fig. 4D). RT-qPCR analysis revealed a significant induction of HIF-1α regulated genes involved in inflammation(59), including cyclooxygenase-2 *(Cox2)* and monocyte chemoattractant protein 1 *(Mcp1)* as well as genes involved in hypoxia(56, 60) metabolic reprograming such as glucose transporter 1 (solute carrier family 2, *Slc2a1*, GLUT-1), lactic acid and pyruvate transporter (solute carrier family 16, *Slc16a3*, also known as monocarboxylate transporter 4 MCT4). In addition, we also observed a significant increase in HIF-1α regulated genes involved in angiogenesis(56, 60), including a cell-surface glycoprotein *Cd44*, Lysyl oxidase-like 2 *(Loxl2)* and angiopoietin-like 4 *(Angptl4)* (Fig. 4D). Importantly, our findings in the CCM animal model were extended to human CCM because we also detected the elevation of HIF 1α regulated genes in human CCM lesions in comparison with lesion-free brain tissue from a CCM patient or noneurological disease control (Fig. 4E). Since we observed that COX-2 is increased at mRNA (Fig. 4D) and protein (Data not shown) levels in *Pdcd10^ECKO^* hindbrains and at the mRNA level in human CCM lesions (Fig. 4E), we next investigated the effect of using celecoxib (40 mg/Kg), a nonsteroidal anti-inflammatory drug (NSAID) that primarily inhibits COX-2 activity, in two preclinical CCM models. Blinded measurements using microCT analysis of neonatal celecoxib-treated *Pdcd10B^ECKO^* mice compared with vehicle-treated littermate *Pdcd10^BECKO^* mice control indicated a reduction in vascular lesion volume principally in the cerebrum (Fig. 5A). Histological analysis of the hippocampal area in P13 *Pdcd10B^ECKO^* mice further demonstrates the high propensity of CCM lesions to develop surrounded by GFAP+ astrocytes, and inhibition of COX-2 reduced the CCM lesions’ density in *Pdcd10^BECKO^* mice (Fig. 5B, fig. S5). Next, we investigated the effect of using celecoxib for 15 days in juvenile *Pdcd10B^ECKO^* mice. Notably, blinded measurements of microCT analysis confirmed reduction in CCM in cerebrum and cerebellum of P80 celecoxib-treated *Pdcd10B^ECKO^* mice (Fig. 5C). Moreover, histological analysis of the cerebellum and hippocampal area in P80 celecoxib-treated *Pdcd10^BECKO^* mice confirmed a decrease in CCM lesions’ density and in GFAP-immunoreactivity (Fig. 5D, fig. S5). Similar to the CCM lesions detected in brain, we observed that P80 *Pdcd10B^ECKO^* mice develop gross CCM lesions through the spinal cord (fig. S6), and with high propensity to develop in GFAP+ astrocytes areas. In addition, P80 *Pdcd10^BECKO^* spinal cords exhibited ~1.6-fold increase in *Vegfa* mRNA levels that it is prevented in P80 celecoxib-treated *Pdcd10B^ECKO^* mice (Fig. 5E), whereas the elevation of Nos3 mRNA remain unchanged (Fig. 5F). Collectively, these results suggest that CCM endothelium can induce HIF-1α dependent hypoxia and angiogenesis programs in astrocytes and that this non-cell-autonomous mechanism contributes to disease. Moreover, the capacity of celecoxib to ameliorate two murine CCM models underscores the potential of therapeutic strategies to emerge from these insights.

**Fig 4.**
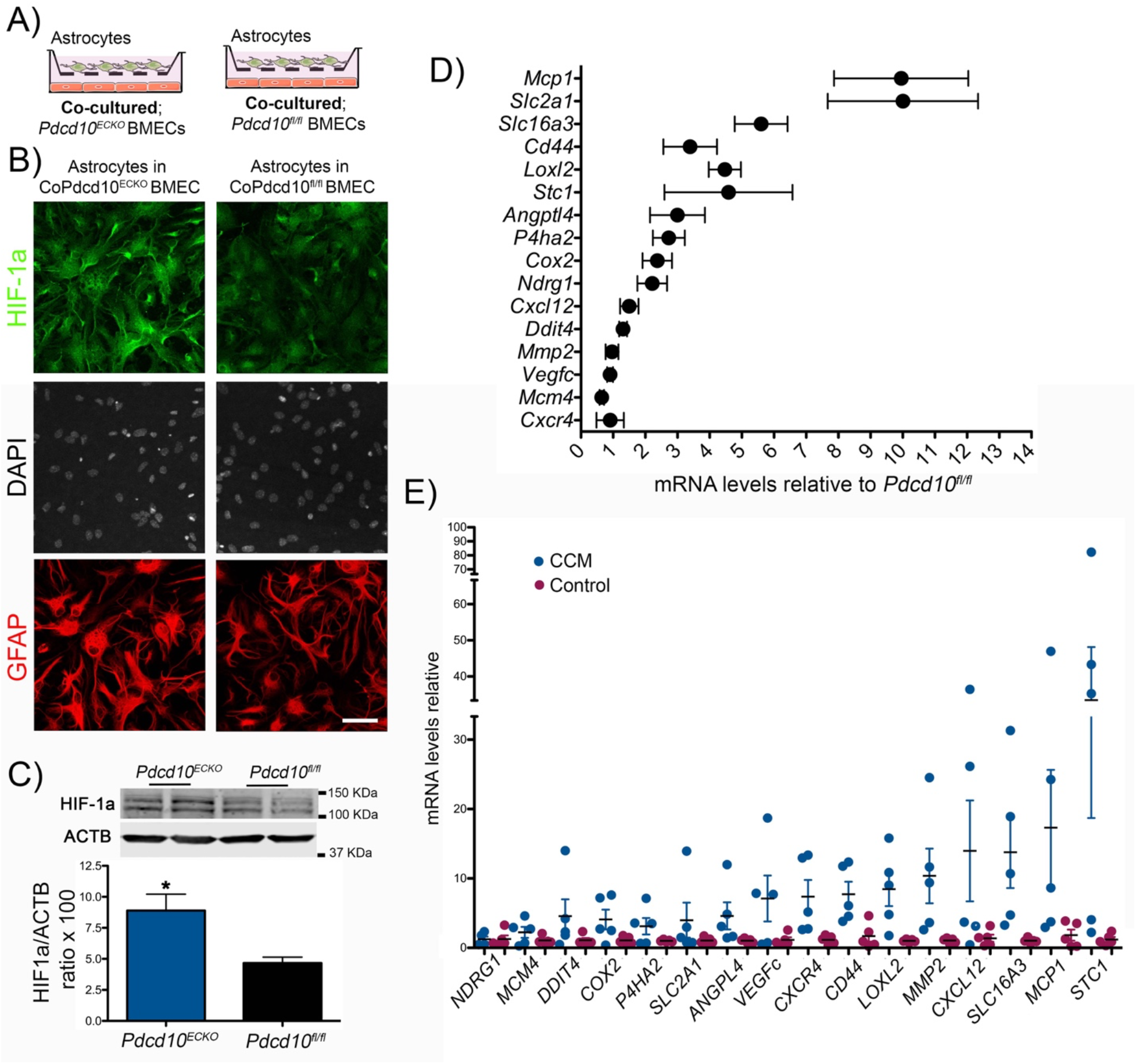
Increase in normoxic HIF-1α stabilization in astrocyte and COX-2 during CCM. (**A**) Schematic diagram of astrocytes co-cultured with *Pdcd10^ECKO^* and *Pdcd10^fl/fl^* BMECs. (**B**) Immunofluorescence staining for HIF-1α (green), GFAP (red), and DAPI for nuclear DNA (white) of primary astrocytes co-cultured with *Pdcd10^ECKO^* BMEC compared to *Pdcd10^fl/fl^* BMEC control for 48hrs (n=3). (**C**) Quantification of HIF-1α in cerebellar tissue in P10 *Pdcd10^ECKO^* control *Pdcd10^0^* littermates, as assessed by Western blot (SEM, n= 4 mice in each group). (**D**) Analysis of HIF-1 a-target genes by RT-qPCR in cerebellar tissue from P10 *Pdcd10^ECKO^* and control *Pdcd10^fl/fl^* littermates (SEM, n=5 or 7 mice in each group). (**E**) Analysis of HIF-1α-target genes by RT-qPCR in human CCM lesions and control human brain tissue (SEM, n=5). Data are mean±SEM. *, P<0.05, determine by Student’s *t* test. Scale bar: (**B**) 50 μm.

**Fig 5.**
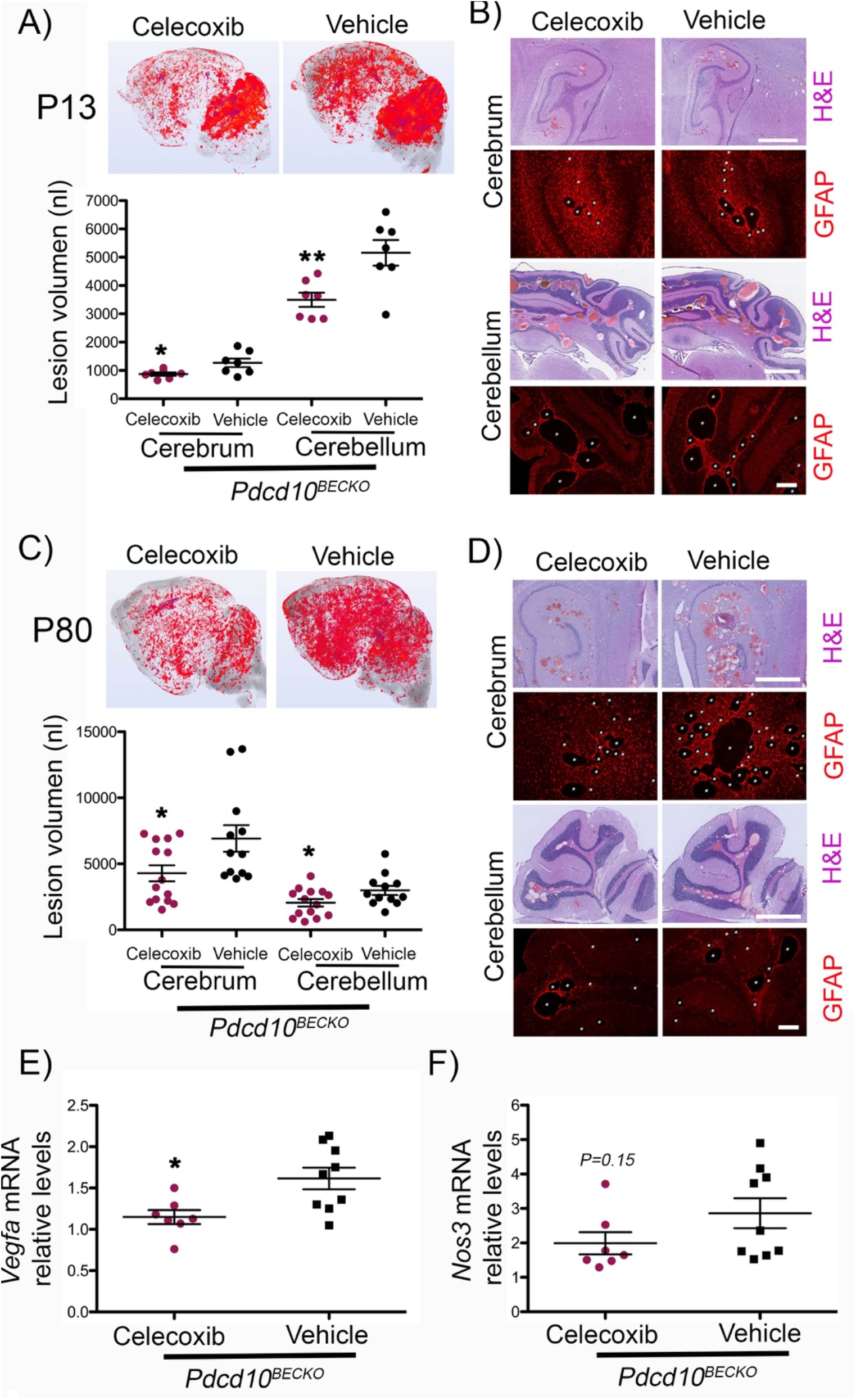
COX-2 inhibition prevent CCM lesions in *Pdcd10^BECKO^* mice. (**A**) Prominent lesions are present in the cerebellum and cerebrum of P13 *Pdcd10B^ECKO^* mice. Intragastric administration of 40 mg/Kg celecoxib for four consecutive days P6 to P9 suppressed lesion formation. Quantification of lesion volumes by micro-CT analysis from mice at P13 treated with celecoxib or vehicle (SEM, n=7 mice in each group). (**B**) Hematoxylin and eosin (pink and purple) or GFAP (red) staining of cerebral (hippocampal area) and cerebellar sections from *Pdcd10^BECKO^* mice after treatment with celecoxib or vehicle (n=3). (**C**) Prominent lesions are present in the cerebellum and cerebrum of P80 *Pdcd10B^ECKO^* mice. Oral gavage administration of 40 mg/Kg celecoxib for fifteen consecutive days P55 to P70 suppressed lesion formation. Quantification of lesion volumes by micro-CT analysis from mice at P80 treated with celecoxib or vehicle (SEM, n=12 or 14 mice in each group). (**D**) Hematoxylin and eosin (pink and purple) or GFAP (red) staining of cerebral (hippocampal area) and cerebellar sections from *Pdcd10^BECKO^* mice after treatment with celecoxib or vehicle (n=3). (**E-F**) Quantification of *Vegfa (**E**)* or Nos3 *(**F**)* mRNA levels in P80 *Pdcd10^BECKO^* spinal cords after treatment with celecoxib or vehicle from experiments in *C* (SEM, n=7 or 9 mice in each group). Data are mean±SEM. *, P<0.05, **, P<0.01; determine by Student’s *t* test. Scale bar: (**B** and **D**) 1 mm (H&E), 200 μm (GFAP).

### Loss of brain endothelial *Pdcd10 or Krit1* increases the expression of eNOS

Previous work has demonstrated that eNOS contributes to VEGF-induced angiogenesis and vascular permeability(41) potentially through stabilization of HIF1-α by Nitric Oxide (NO). Genetic inactivation of *Pdcd10 (Pdcd10^ECKO^)* or *Krit1 (Krit1^ECKO^)* results in increased *Nos3* mRNA levels(6, 61). When we co-cultured *Pdcd10^ECKO^* endothelial cells with astrocytes we observed a dramatic upregulation of *Nos3* mRNA (~19.2 fold increase) associated with increased eNOS protein expression (~8.9 fold increase) in *Pdcd10^ECKO^* BMECs when compared to *Pdcd10^fl/fl^* BMEC control as assessed by Western blot analysis (fig. S7). Similar results were observed in *Krit1^ECKO^* BMECs: a ~7.4 fold increase *Nos3* mRNA was associated with increased eNOS protein expression (~1.9 fold increase) (fig. S7). These results indicate that upon genetic inactivation of *Pdcd10* or *Krit1* in BMECs, there is a significant increase in eNOS mRNA and protein.

To investigate the distribution and expression of eNOS during CCMs in vivo, we first used neonatal *Pdcd10^ECKO^* mice. Consistent with results observed in vitro, *Pdcd10^ECKO^* hindbrains showed ~3.3 fold increase in *Nos3* mRNA levels compared with littermate *Pdcd10^fl/fl^* controls (Fig. 6A). Furthermore, Western blot analysis showed a ~5.8 fold increase in eNOS protein expression in *Pdcd10^ECKO^* hindbrains relative to *Pdcd10^fl/fl^* hindbrains (Fig. 6B). Immunohistochemistry analysis of hindbrain sections revealed that eNOS was upregulated in CCM lesions. As showed in Fig. 6C, we observed that eNOS staining was increased in *Pdcd10^ECKO^* dilated vasculature as showed by colocalization of antibodies specific against eNOS and isolectin B4-FITC staining, which specifically labels the brain vasculature (Fig. 6C). Moreover, we analysis of the distribution and expression of eNOS in juvenile CCM animal model indicated that *Nos3* mRNA levels were elevated in brain tissue in *Pdcd10^ECKO^* mice (Fig. 6D,E). We also observed robust and focal upregulation of eNOS protein in the lesions of *Pdcd10^ECKO^* mice (Fig. 6F). Together, these results demonstrate that eNOS mRNA and protein expression is increased in CCM lesions.

**Fig 6.**
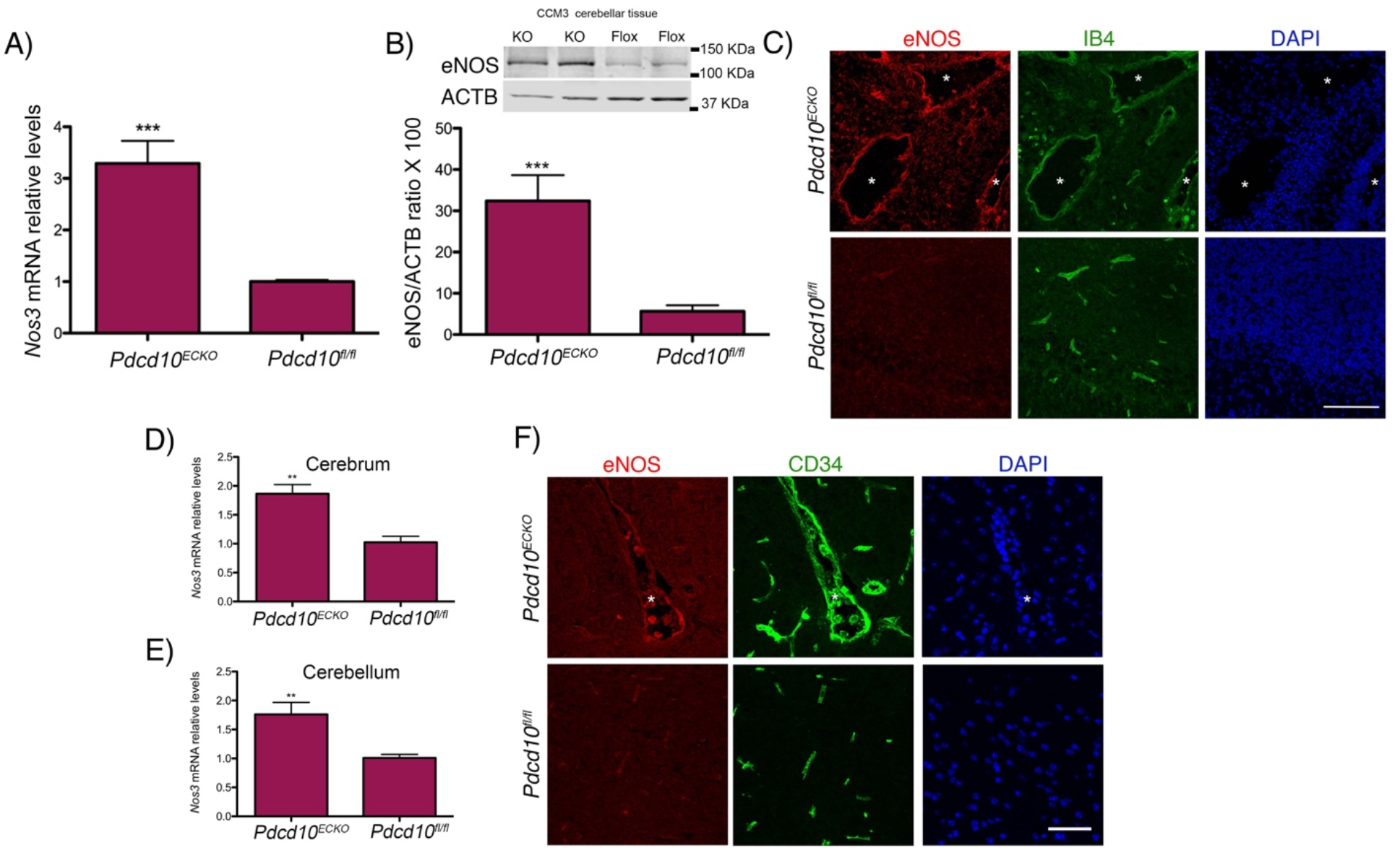
Loss of brain endothelial *Pdcd10* increases the expression of eNOS in situ. (**A**) Analysis of *Nos3* mRNA levels by RT-qPCR in hindbrains of P10 *Pdcd10^ECKO^* and littermate *Pdcd10^fl/fl^* control (SEM, *n*=4 or 5 mice in each group). (**B**) Analysis of eNOS levels in hindbrains of P10 *Pdcd10^ECKO^* and littermate *Pdcd10^fl/fl^* control, as assessed by Western blot analysis (SEM, *n*=4 or 5 mice in each group). (**C**) Confocal microscopy of cerebellar cortex P10 stained for eNOS (red), endothelial marker isolectin B4 (green), and DAPI for nuclear DNA (blue). Asterisks indicate vascular lumen of CCM lesions (*n*=4 mice in each group). (**D**-**E**) Analysis of *Nos3* mRNA levels by RT-qPCR in brains of P30 *Pdcd10^ECKO^* and littermate *Pdcd10^fl/fl^* control. (**F**) Immunofluorescence staining of eNOS (red), endothelial marker cd34 (green), and DAPI for nuclear DNA (blue) (*n*=4 or 6 mice in each group). Data are mean±SEM. **, P<0.01, ***, P<0.001; determine by Student’s *t* test. Scale bars: (**C**) 100 μm; (**F**) 50 μm.

### Increased eNOS expression in human CCMs is regulated by KLF2 and KLF4

In human CCM brain tissue, there was a ~3 fold increase in *NOS3* mRNA levels in comparison to lesion-free brain tissue (Fig. 7A). We also observed that eNOS staining was increased in human CCM endothelium in comparison with lesion-free brain tissue (Fig. 7B). Transcription factors KLF2 and KLF4 are regulators of eNOS(62, 63). Therefore, we confirmed that the changes in brain endothelial eNOS, at the protein and mRNA levels, were associated with an increase in transcription factors KLF2 and KLF4 (Fig. 7C,D). Moreover, we observed that reducing the expression of KLF2 and KLF4(37) prevented increased eNOS expression in KRIT1-depleted human brain endothelial cells (Fig. 7E). These results suggest that the increase in KLF2 and KLF4 increased brain endothelial eNOS in CCMs (Fig. 7F).

**Fig 7.**
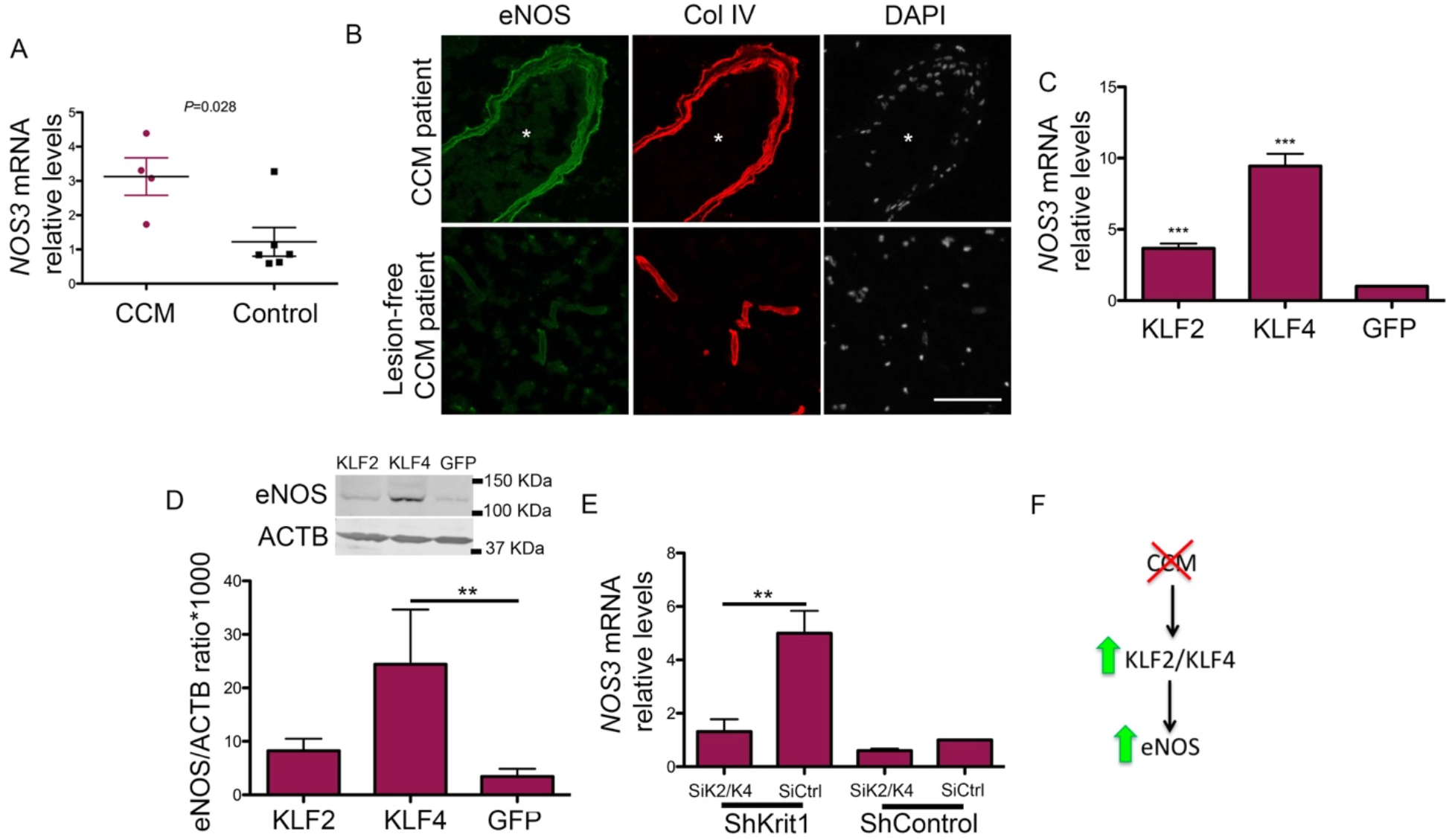
KLF2 and KLF4 regulate increased eNOS expression during CCM. (**A**) Expression levels of *NOS3* mRNA as assessed by RT-qPCR from human CCM lesions and compared to non-neurological disease control (SEM, n=4 or 6 in each group). (**B**) Immunofluorescence staining of eNOS (green) and collagen IV (Col IV; red) of human CCM lesion matched to CCM lesion-free brain tissue (n=3). Asterisks denotate vascular lumen of CCM lesion. Nuclei were counterstained with DAPI (white). (**C**) Human umbilical vein endothelial cells (HUVECs) were transduced with lentivirus encoding KLF2 or KLF4, as previously reported(6), and analysis of *NOS3* mRNA levels by RT-qPCR were determined in cells overexpressing KLF2 or KLF4 and compared with lentivirus encoding GFP as control (n=3 or 4). (**D**) Analysis of eNOS protein levels in HUVECs transduced with lentivirus encoding KLF2 or KLF4, as determined by Western blot analysis(37); lentivirus encoding GFP was used as control (SEM, n=3). (**E**) Analysis of *NOS3* mRNA levels by RT-qPCR in hCMEC/D3 cells transduced with lentivirus encoding shKRIT1 or scrambled control, followed by transfection with KLF2 and KLF4-specific small interfering RNAs (siRNA;siK2/K4) or small interfering RNA control (siCtrl) (SEM, n=4). (**F**) Schematic model. KLF2 and KLF4-mediated elevation of eNOS in CCM endothelium by a cell-autonomous mechanism. Data are mean±SEM. **, P<0.01, ***, P<0.001; determine by Student’s *t* test and 1-way ANOVA, followed by the Tukey post hoc test. Scale bar:(B) 100 μm.

### Loss of brain endothelial *Pdcd10* increases nitric oxide production via eNOS

Endothelial cells metabolize L-arginine via eNOS to produce nitric oxide (NO), therefore we next investigated if eNOS upregulation increases NO production in *Pdcd10^ECKO^* BMEC. Using a colorimetric assay to quantitatively measure total NO2 and NO3 (nitrites and nitrates) in the culture media, we observed that *Pdcd10^ECKO^* BMEC possessed a ~1.7 fold increase in NO release when compared with *Pdcd10^fl/fl^* BMEC control (NO basal levels: ~3 μM) (Fig. 8A). The increase in NO release is ascribable in part to the upregulation of eNOS, because genetic inactivation in one copy of the *Nos3* gene *(Nos3^+/-^)* significantly reduced NO production in *Pdcd10^ECKO^* BMEC (Fig. 8B). Consistent with these results, low levels of *Nos3* mRNA (data not shown) and eNOS protein in *Pdcd10^ECKO^;Nos3^+/-^* BMEC were observed (Fig. 8C). Moreover, addition of a nitric oxide synthase (NOS) inhibitor, L-NAME, reduced NO production (Fig. 8B) without changing eNOS mRNA and protein levels (data not shown). Taken together, these studies show that the loss of *Pdcd10* in BMECs leads to an increase in eNOS expression and subsequent endothelial NO release.

**Fig 8.**
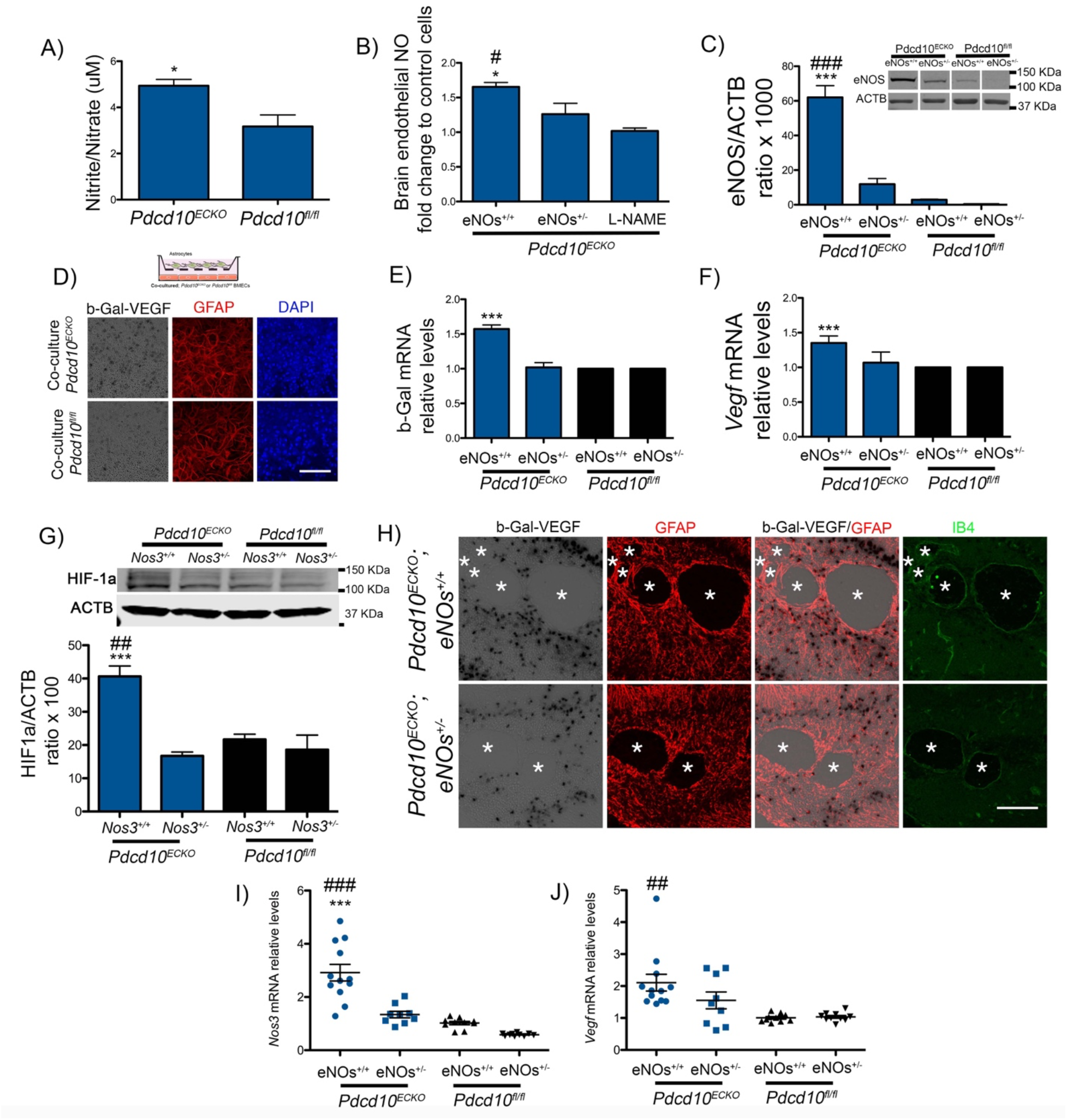
Loss of brain endothelial *Pdcd10* increases nitric oxide production and induces astrocyte-derived VEGF. (**A**) Total nitrite and nitrate concentration as an intermediate of NO production assessed by colorimetric assay analysis from the media of *Pdcd10^ECKO^* and *Pdcd10^fl/fl^* BMEC cultured for 36hrs (SEM, n = 7). (**B**) Increased NO release in *Pdcd10^ECKO^* is significantly reduced in *Pdcd10^ECKO^;Nos3+/-* BMEC or following incubation with a NOS inhibitor, L-NAME (150μM) (SEM, n = 3). (**C**) Quantification of eNOS protein from *Pdcd10^ECKO^* and *Pdcd10^ECKO^ Nos3^+/-^* BMECS relative to *Pdcd10^fl/fl^*and *Pdcd10^fl/fl^;Nos3^+/-^* BMECs controls (SEM, n = 5). Culture media was supplemented with 500μM L-Arginine and was deficient in serum. Lanes in this panel were run on the same gel but were noncontiguous. (**D**) β-gal/VEGF expression, as shown by X-gal staining (black) and immunofluorescence staining for GFAP (red) of primary cultured astrocytes co-cultured with *Pdcd10^ECKO^* BMEC compared to *Pdcd10^fl/fl^* BMEC control for 48hrs (n=2). (**E**) RT-qPCR analysis of *β-gal* and (**F**) *Vegfa* mRNA in primary cultured astrocytes cocultured with *Pdcd10^ECKO^* BMEC compared to *Pdcd10^fl/fl^* BMEC control for 24hrs (SEM, n = 4). (**G**) Quantification of HIF-1α protein from primary astrocytes co-cultured with *Pdcd10^ECKO^* BMEC and *Pdcd10^ECKO^;Nos3+/-* BMEC relative to astrocyte co-cultured with *Pdcd10^fl/fl^* or *Pdcd10^fl/fl^;Nos3^+/-^* BMECs controls for 48hrs (SEM, n = 4). (**H**) Confocal microscopy of neonatal hindbrain at P10 from *Pdcd10^ECKO^;Nos3^+/+^; Vegfa^tm1^-^1Nagy^ (Pdcd10^ECKO^;eNOS^+/+^)* and *Pdcd10^fl/fl^;Nos3^+/-^;Vegfa^tm1.1Nagy^ (Pdcd10^ECKO^;eNOS^+/-^)* littermate control stained for GFAP-positive astrocytes (red), β-gal/VEGF expression detected by X-gal staining (black), isolectin B4 (green). Asterisks, vascular lumen of CCM lesions (n=3). (**I-J**) Quantification of *Nos3 (**I**)* and *Vegf (**J**)* mRNA levels in P10 *Pdcd10^ECKO^;Nos3^+/-^* and *Pdcd10^ECKO^* and *Pdcd10^fl/fl^;Nos3^+/-^* hindbrains when compared with littermate *Pdcd10^fl/fl^* controls, as assessed by RT-qPCR (SEM, n=9 or 12 mice in each group). Data are mean±SEM. ^*,#^, P<0.05, ^##^, P<0.01, ***,###, P<0.001 (* comparison to *Pdcd10^ECKO^;Nos3^+/-^* and # comparison to L-NAME or *Pdcd10^fl/fl^;eNOs^+/+^*); determine by Student’s *t* test and 1-way ANOVA, followed by the Tukey post hoc test. Scale bars: (D and H) 100μm.

### Brain endothelial nitric oxide induces astrocyte-derived VEGF following loss of *Pdcd10*

We showed evidence of increased levels in VEGF expression in GFAP+ brain and retinal astrocytes in *Pdcd10^ECKO^* mice (Fig. 2A,C,D,3A,B, fig. S2,3). However, our in vivo experiments did not unequivocally exclude expression of VEGF in other CNS-resident cells during CCMs, including neurons, microglia and pericytes. Thus, we complemented those studies by performing co-culture experiments between CCM endothelium and astrocytes. First, NO donors, such as DETANONOate, promote brain angiogenesis by upregulation of VEGF signaling(43). To investigate whether NO stimulates the expression of VEGF in astrocytes during CCMs, we prepared primary mouse astrocyte cultures from *Vegfa^tm1.1Nagy^* mice (Fig. 8D). We observed that the purity of cultured astrocytes was high as determined by cells that are double positive for specific astrocyte markers GFAP and integrin B5 (fig. S4). We also observed that our primary astrocyte cultures respond to exogenous NO (0.5mM DetaNONOate (NO donor)) and that elevated levels of NO can increase astrocyte-derived VEGF (as assessed by increase in β-gal/VEGF expression) (fig. S8). We next investigated whether elevated brain endothelial NO in *Pdcd10^ECKO^* results in increased astrocyte-derived VEGF levels. Therefore, we co-cultured purified astrocytes in the presence of *Pdcd10^ECKO^* or *Pdcd10^fl/fl^* BMECs in serum-free conditions supplemented with L-arginine, and measured changes in β-gal/VEGF expression in astrocytes. We observed that *Pdcd10^ECKO^* BMEC significantly increase β-gal/VEGF expression in astrocytes in a co-culture system (Fig. 8D,E). These results were consistent with increased levels of β-gal and *Vegfa* mRNA levels in astrocytes co-cultured with *Pdcd10^ECKO^* BMEC compared to cocultured with *Pdcd10^fl/fl^* BMEC control (Fig. 8E,F). Furthermore, genetic inactivation of one copy of the *Nos3* gene in *Pdcd10^ECKO^* BMEC was sufficient to prevent stability of HIF-1α and subsequent *Vegfa/*β-gal upregulation in astrocytes indicating that the increase in astrocyte-derived *Vegfa/*β-gal mRNA levels were specific to the upregulation of eNOS (Fig. 8E,F) and dependent of HIF-1α stability (Fig. 8G).

We next investigated whether the increase in astrocyte-derived VEGF during CCMs was associated with elevation of eNOS in mice. *Vegfa^tm1.1Nagy^* mice were intercrossed with CCM animal models and with animals deficient in one copy of the *Nos3* gene (eNOs+/-). We observed that CCM lesions were slightly reduced and with a partial decrease in β-gal/VEGF expression in P10 *Pdcd10^ECKO^;Vegfa^tm1.1Nagy^;Nos3^+/-^* hindbrains when compared with *Pdcd10^ECKO^; Vegfa^tm1.1Nagy^*;Nos3^+/+^ littermate control (Fig. 8H). However, the strong induction of *Nos3* expression following loss of brain endothelial *Pdcd10* (Fig. 8I) suggest that a substantial level of eNOS/NO signaling could contribute to partial elevation in VEGF expression in *Pdcd10^ECKO^; Nos3^+/-^* hindbrains (Fig. 8J) and an incomplete rescue (Data not shown). Altogether our data provide key insights into understanding a circuit of neurovascular dysfunction during CCM disease mediated between the CCM endothelium and astrocytes (Fig. 9). These data provide strong evidence that the nitric oxide (NO) production as a consequence of the elevation of eNOS in CCM endothelium, through elevation of KLF2/KLF4, contributes to the normoxic stability of HIF-1α and subsequent elevation in astrocyte-derived VEGF. Our findings further support that the hypoxic program components, such as COX-2, represent potential therapeutic targets for CCMs disease (Fig. 9).

**Fig 9.**
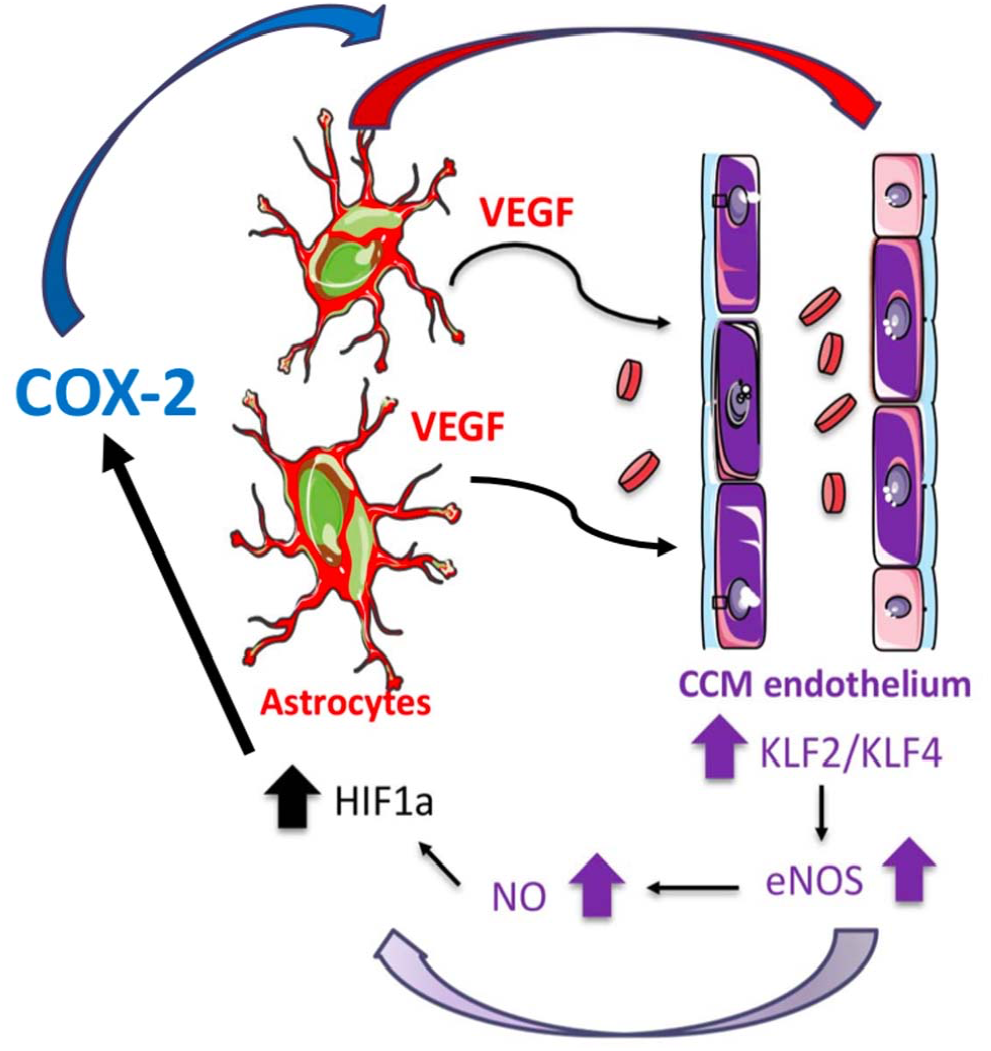
Astrocytes integrate a circuit of neurovascular dysfunction during CCM disease. Model by which astrocytes make a significant contribution to CCM pathogenesis. An increase in astrocyte VEGF synthesis is driven by endothelial nitric oxide (NO) generated due to KLF2 and KLF4-dependent elevation of eNOS in CCM endothelium. Production of NO in CCM endothelium stabilizes HIF-1α in astrocytes, resulting in increased VEGF production and expression of a “hypoxic” program under normoxic conditions. Pharmacological inhibition of HIF1-driven COX-2 can ameliorate murine models of CCM disease.

## Discussion

The propensity of CCM lesions to form in the CNS parenchyma has never been mechanistically clarified. Our study shows that astrocytes respond to CCM endothelium to initiate a non-cell-autonomous enhancement of CCM formation mediated by astrocyte hypoxia and angiogenesis programs. Our experiments show that elevation of eNOS in CCM endothelium contributes to increase in astroglia-derived VEGF and HIF-1α protein stabilization in astrocytes during CCM disease. The increase in eNOS expression was ascribable to an elevation of the KLF2 and KLF4 transcription factors in CCM. eNOS production in the CCM endothelium results in a sustained elevation of NO that leads to normoxic HIF-1α protein stabilization and increase in HIF activity in CCM tissue. We therefore propose that astrocytes contribute to CCM disease by a non-cell-autonomous mechanism mediated by CCM endothelium-driven hypoxia and angiogenic programs.

Our in vivo observations in the neonatal CCM animal model, using endothelial-specific inactivation of *Pdcd10* or *Krit1*, revealed that lesion development is spatially restricted to zones of fibrous astrocytes, which are prevalent in the white matter of the CNS(46), and that the selective ablation of proliferative astrocytes(48) markedly reduced CCM lesion and an increase in VEGF expression, prompting the suggestion that a subset of proliferative astrocytes supports CCM lesions in the developing hindbrain. Moreover, our observations in the chronic CCM animal model(53, 54), using brain endothelial-specific inactivation of *Pdcd10*(64), further revealed the high propensity of CCM lesions to develop surrounded by GFAP+ astrocytes throughout the CNS, including the cerebellum, cerebrum, and spinal cord. Earlier reports have shown that loss of *Pdcd10* in neuroglia cells leads to formation of CCM lesions through non-cell-autonomous mechanisms, implicating a role of neural cells in the CCM pathogenesis(17). Consistent with these findings, astrogliosis has been shown to intermingle with granulation tissue in human CCM(65). However, while some subsets of reactive and proliferative astrocytes can be detrimental and contribute to CNS pathologies while other subsets of reactive astrocytes can be beneficial by supporting CNS recovery (24, 48, 66, 67). The mechanisms that contribute to functional heterogeneity in reactive astrocytes are not completely understood. Recent studies have shown that microglia control astrocyte pathogenic activities during neuroinflammation by releasing of VEGF-B and TGFalpha (68), and during neurodegeneration by secreting of Il-1alpha, TNF and C1q (24). Our findings revealed that the CCM endothelium controls astrocyte responses through increasing the levels of eNOS and subsequent release of brain endothelial NO. Vascular eNOS has important roles in regulating endothelial homeostasis by producing tonic NO levels, but upregulation of the eNOS gene or activated form has been previously implicated in vascular permeability, angiogenesis and neuroinflammation(26, 42, 43, 69). Here we show that eNOS is increased in human CCM endothelium and the upregulation in endothelial KLF2 and KLF4 can account for the augmented eNOS expression at the transcriptional level (63). However, these studies do not exclude the possibility that the elevation of endothelial VEGF signaling during CCM(6) may also contribute to a sustained eNOS activation (41, 70) by posttranslational modification via PI3K/AKT, Ca^2+^/calmodulin signaling (71) during long-term adaptations, and this notion needs to be further tested.

CCMs are hypersensitive to angiogenesis due to increase in VEGF signaling, the loss of an anti-angiogenic checkpoint protein, TSP1 (6, 38), augmented secretion of angiopoietin-2(11), and deregulation of Notch signaling(72). The increase in VEGF signaling maybe associated with elevated levels of angiogenic factor VEGF in CCM lesions of individuals with the hereditary and sporadic form of the disease(29–32). We found that CCM endothelium-induced elevation of astrocyte-derived VEGF in neonatal and juvenile brains, and elevation of VEGF levels are dependent on eNOS signaling. Notably, an increase in astrocyte-derived VEGF can contribute to the development and progression of CNS disease (26–28) by disassembly of interendothelial junctions(6, 73, 74). Importantly, early studies show that eNOS contributes to VEGF-induced angiogenesis through the intercellular messenger NO (41–43, 75). Furthermore, it has also been shown that NO-mediated astrocyte HIF-1α stabilization modulates transcriptional and metabolic activities(75, 76). In agreement with this, we observed elevated levels of eNOS protein and NO bioavailability in *Pdcd10^ECKO^* BMEC that increased levels of HIF-1α in co-cultured astrocytes. Loss of one copy of the gene *Nos3* in *Pdcd10^ECKO^* BMEC is sufficient to attenuate astrocytes HIF-1α stabilization in the co-culture system. Importantly, although we found that astrocytes respond to CCM endothelium, other cells of the neurovascular unit (e.g., neurons, pericytes, and microglia) may also contribute to CCM disease in a non-cell-autonomous manner through normoxic stabilization of HIF-1α protein. It is tempting to speculate that CCM endothelium may harmonize cell activation in the neurovascular unit by elevation of NO during CCM disease. Because, NO can reach cells millimeters away from where it is produced, and this in turn can stabilize HIF-1α protein(77, 78) from degradation by several mechanisms; directly by HIF-1α nitrosylation or also indirectly by increasing production of reactive oxidative species or by binding to the iron co-factor required for prolyl hydroxylases activity(79). HIF-1α can translocate to the nucleus where it dimerizes with HIF-1β and binds to the hypoxia-responsive element in the promoter region of several target genes(60). Our data indicate that in mouse and human CCM tissue there is an activation of the HIF-1α program associated with the glycolytic pathway by increasing GLUT-1 that guarantees rapid energy production, and an increase in MCT4 that promotes efficient removal of lactic acid, an end product of glycolysis metabolism(60). Elevated HIF activity in CCM tissue was also associated with increases in several pro-angiogenic and inflammatory factors produced by glial cells, including ANGPTL4 that can induce vascular dysfunction in CNS pathologies by triggering angiogenesis(80, 81) and COX-2 an enzyme that catalyzes the biosynthesis of prostanoids during inflammation(59). We focused on studying COX-2 inhibition during CCM because it has been shown that COX-2 inhibition can suppress angiogenesis in vitro and in vivo (82, 83). Moreover, selective COX-2 inhibitors are safe and well-tolerated drugs that can be repurposed for therapy for CCM disease. Our results show that pharmacological inhibition of COX-2 significantly reduced vascular lesion formation in two models of CCM disease. Importantly, we notice that the inhibition of COX-2 in the chronic CCM animal model results not only in a decrease in CCM lesions’ density but also in GFAP-immunoreactivity, suggesting that amelioration of lesion genesis counterparts astrocyte activity in the lesions. In line with these results, a recent retrospective cohort study reported that the use of NSAIDs was correlated with lower risk of hemorrhage among patients affected by CCMs(84). However, this uncontrolled association cannot be interpreted as proof of therapeutic benefit, as patients with recent CCM hemorrhages were inherently less likely to be taking NSAIDs. Cox-2 inhibitors must be investigated for safety and effectiveness in prospective controlled trials, and specific dose-effect and duration of treatment must be carefully defined. A platform for trial readiness, exploring proof of concept effect on lesional bleeding in human subjects, is currently being developed, and can efficiently be applied to repurposed drugs, like NSAIDs(85). The precise role of NSAIDs in reducing CCM risk of hemorrhage version lesion formation remains incompletely understood. The COX-2-prostaglandin pathway has been implicated in vascular sprouting, migration, and induction of growth factors and proteolytic enzymes that alter angiogenesis(82, 83, 86, 87), suggesting that COX-2 could be an important therapeutic target for pathological angiogenesis(82). Moreover, inhibition of COX-2 augment VEGF pathway blockade during refractory tumor angiogenesis(88), suggests that combinatorial COX-2/VEGF pathway inhibition could be used as a potential approach to ameliorate/prevent CCMs. In addition, Sulindac metabolites, a NSAID, has been shown to inhibit b-catenin-stimulate transcription of endothelial-mesenchymal transition markers and development of CCMs(89). However, additional studies will be required to determine whether COX-2 inhibitors prevent an increase in astroglia-derived VEGF in response to CCM endothelium. Collectively our data are consistent with the notion that reciprocal communication between CCM endothelium and astrocytes drive CCM lesion formation and contribute to neurovascular dysfunction. These observations point to the possibility of designing therapeutic approaches aimed at preventing endothelial dysfunction and astroglia activation as an intervention to reduce the burden of CCM disease in humans.

## Materials and Methods

### Human tissue and genetically modified mice

Human brain samples of patients with CCM and control without neurological disease were obtained from the Angioma Alliance Tissue Bank. Endothelial-specific conditional *Pdcd10-null* mice were generated by crossing a *Pdgfb* promoter-driven tamoxifen-regulated Cre recombinase *(iCreERT2)*(90) with loxP-flanked *Pdcd10 (Pdcd10 ^fl/fl^* generous gift from Wang Min, Yale University; *Pdgfb-iCreERT2;Pdcd10^fl/fl^)* mice. Brain endothelial-specific conditional *Pdcd10*-null mice were generated by crossing a *Slco1c1* promoter-driven tamoxifen-regulated Cre recombinase(64) *(iCreERT2*, generous gift from Markus Schwaninger) with *Pdcd10 ^fl/fl^* mice. eNOS-deficient mice *(Nos3^-/-^)* were obtained from Jackson Laboratory and crossed with *Pdgfb-iCreERT2;Pdcd10^fl/fl^* mice to generate *Pdgfb-iCreERT2; Pdcd10^fl/fl^;Nos3^-/-^* and *Pdgfb-iCreERT2; Pdcd10^fl/fl^;Nos^+/-^* mice. On postnatal day 3, mice were administered 50 μg of 4-hydroxi-tamoxifen [H7904; Sigma-Aldrich] by intragastric injection to induce genetic inactivation of the endothelial *Pdcd10* gene in littermates with *Pdgfb-iCreERT2 (Pdcd10^ECKO^)*, and *Pdcd10^fl/fl^* mice were used as littermate controls. For CCM in juvenile animals, On postnatal day 6, mice were administered 100 μg of 4-hydroxi-tamoxifen [H7904; Sigma-Aldrich] by intragastric injection to induce genetic inactivation of the endothelial *Pdcd10* gene in littermates with *Pdgfb-iCreERT2 (Pdcd10^ECKO^)*, and *Pdcd10^fl/fl^* mice were used as littermate controls. On postnatal day 1, mice were administered 50 μg of tamoxifen [H7904; Sigma-Aldrich] by intragastric injection to induce genetic inactivation of endothelial *Pdcd10* gene in littermates with *Slco1c1-iCreERT2; (Pdcd10B^ECKO^)*, and *Pdcd10^fl/fl^* mice were used as littermate controls. *Vegfa^tm1.1Nagy^* mice, expressing a B-galactosidase (LacZ) reporter gene inserted into the 3′ untranslated region of the *Vegfa* gene(50), were obtained from the Jackson Laboratory. The *GFAP-TK* mice line was used to selectively ablate proliferative astrocytes *(Glial fibrillary acidic protein-Thymidine kinase*, generous gift from Michael V. Sofroniew)(47, 48).

For celecoxib treatment, Brain endothelial-specific conditional *Pdcd10-null* mice were randomized (by flipping a coin) to celecoxib (40 mg/Kg, celebrex) or vehicle treatment (0.5% methylcellulose plus 0.025% tween20). For neonatal experiments, 100 μl celecoxib or vehicle was administered by intragastric injection on P6,P7,P8,P9 and animals were sacrifice at P13. For adult experiments, males and females brain endothelial-specific conditional *Pdcd10*-null mice were randomized (by flipping a coin) to celecoxib (40 mg/Kg, celebrex) or vehicle treatment (0.5% methylcellulose plus 0.025% tween20). Celecoxib or vehicle was administered by oral gavage administration for fifteen consecutive days P55 to P70 and animals were sacrifice at P80.

### Isolation of primary astrocytes

*Vegfa^tm1.1Nagy^* mice or *Hif-1α^fl/fl^* mice with transgenic mice expressing the tamoxifen-inducible recombinase CreERT2 under the control of the astrocyte *Aldh1l1* promoter at postnatal day 5-7 were sacrificed, and their brains were isolated and placed into cold solution A (0.5% bovine serum albumin (BSA) in DMEM and 1 μg/μl glucose, 10mM HEPES, 1x penicillin-streptomycin). Brain cortices were separated from the brain and rolled on dry filter paper to detach and remove the meninges. Cortices from 8-11 mice were pooled and minced with scissors in solution A, and the tissue was centrifuged at 215g for 5 minutes at 4°C. The tissue pellet was digested with a papain solution (0.7mg/ml papain suspension [LS003126; Worthington], 20units/ml DNase I [11284932001; Sigma-Aldrich], and 0.150μg/ml tosyl-lysine-chloromethyl-ketone [T7254; Sigma-Aldrich]) at 37°C for 25 min with vigorous shaking every 10min. The tissue suspension was triturated using thin-tipped Pasteur pipettes until partially homogenous and centrifuged at 215g for 5 minutes. The pellet was resuspended with solution B (25% BSA in DMEM and 1 μg/μl glucose, 10mM HEPES, 1x penicillin-streptomycin) and centrifuged at 1000g for 20min at 4°C. The lighter phase containing astrocytes was extracted, resuspended in 50ml of solution C (DMEM-1 μg/μl glucose, 10mM HEPES, 1x penicillin-streptomycin), and centrifuged at 215g for 10 minutes at 4°C. The pellet was resuspended again in 50 ml of solution C and centrifugation was repeated.

### Astrocyte culture conditions

The purified primary astrocytes were plated on a Poly-L lysine-coated plate cultured in astrocyte media comprised of 1:1 Neurobasal media and DMEM (1 μg/μl glucose) supplemented with the following: 0.1mg/ml BSA, 0.1 mg/ml transferrin, 0.016mg/ml putrescine, 0.025μg/ml progesterone, 0.016μg/ml sodium selenite, 5ng/ml H-BEGF, 5μg/ml N-acetyl cysteine, 1mM sodium pyruvate, 1x penicillin-streptomycin, and 292μg/ml L-glutamine(24). The primary astrocyte culture identity and purity were confirmed by GFAP and integrin β5 immunofluorescence.

### Growth surface preparation

Poly-L lysine (P8920-100ml, 0.1% (w/v) in H20, Sigma-Aldrich) stock solution was diluted 1 in 10 in Hank’s balanced salts solution plus calcium (HBSS+Ca) (14025092, Sigma-Aldrich) and left for 1h at 37°C on the plastic surface of 6-well plate format. Collagen type I (C8919, 0.1% (w/v) in 0.1 M acetic acid) stock solution was diluted 1 in 20 in HBSS+Ca and left for 1h at room temperature (RT) on the plastic surface of 6-well plate format. For experiments that used transwell polyester membrane inserts (0.4 μm pore, CLS3450 24mm or CLS3460 12mm diameter, Corning Costar), the filters were first coated with Poly-L lysine as described. Coating solutions were removed, and cells were seeded onto the plastic surface or inserts.

### Isolation of primary brain microvasculature endothelial cells

Adult mice 2-4 months old were sacrificed, and brains were isolated and placed into cold solution A. Meninges and choroid plexus were detached and removed, and the brains of 5-6 mice were pooled together and minced with scissors in solution A. Brain tissue suspension was centrifuged at 215g for 5 minutes at 4°C. The tissue was digested with a collagenase/dispase solution (1mg/ml collagenase/dispase [10269638001; Sigma-Aldrich], 20 units/ml DNase I [11284932001; Sigma-Aldrich], and 0.150μg/ml tosyl-lysine-chloromethyl-ketone [T7254; Sigma-Aldrich] in DMEM]) at 37°C for 1h with vigorous shaking every 10min. Then the tissue suspension was triturated using thin-tipped Pasteur pipettes until fully homogenous and centrifuged at 215g for 5 minutes. The pellet was resuspended in cold solution B and centrifuged at 1000g for 20min at 4°C. The lighter phase was discarded and the heavy phase containing the brain microvasculature was digested in collagenase/dispase a second time for 30min at 37°C with vigorous shaking every 10min. After incubation, the suspension was centrifuged (215g for 5min at 4°C) and the pellet was resuspended in BMEC-media that comprised of EBM-2 medium (Lonza) supplemented with the following: 0.025% recombinant human EGF, 0.1% insulin-like growth factor, 0.1% gentamicin, 0.04% ascorbic acid, 0.04% hydrocortisone, and 20% FBS. The BMECs were plated in collagen-coated wells (0.005% collagen in HBSS [C8919, Sigma-Aldrich]) and cultured in 10μg/ml of puromycin for 2 days, followed by 2μg/ml for 2 days(91). Primary BMEC culture identity and purity were confirmed by RNA expression levels of endothelial-specific genes, morphology, and immunofluorescence.

### Inactivation of *Krit1* or *Pdcd10* gene in primary BMECs

After 5 days in culture at 37°C in 95% air and 5% CO2, primary BMEC from mice bearing *Pdcd10^fl/fl^* or *Pdgfb-iCreERT2;Pdcd10^fl/fl^* were passaged to equal confluency (~ 2.5 × 10^5^ cells) on collagen-coated 6-well plates. On day 6-7 from initial culture, *Pdgfb-iCreERT2;Pdcd10^fl/fl^* BMECs were treated with 5μM of 4-hydroxy-tamoxifen (H7904; Sigma-Aldrich) for 48h to delete *Pdcd10 (Pdcd10^ECKO^). Pdcd10^fl/fl^* BMECs were also treated with 4-hydroxy-tamoxifen and used as a control. The deletion of *Pdcd10* in *Pdcd10^ECKO^* BMECs was verified by RT-qPCR analysis. The medium was replaced with fresh BMEC-media (2.5 Isolation of primary brain microvasculature endothelial cells) and changed again every two days(91).

### Co-culture of BMECs and astrocytes

*Pdcd10^ECKO^* and *Pdcd10^fl/fl^* BMECs at passage 1-3 were plated on collagen-coated 6-well plates and maintained in BMEC-media for 15-20 days, while mouse primary astrocytes (~3.5 × 10^5^ cells) were seeded on poly-L-lysine coated transwell filters (3450; Sigma-Aldrich) and maintained in astrocyte-media. Astrocytes were maintained for 3 days before transwells were placed onto the BMEC wells containing astrocyte media supplemented with 500μM L-arginine. BMEC and astrocytes were maintained in co-culture for the time indicated in each experiment.

### Immunofluorescence microscopy

Astrocytes were grown on poly-L-lysine coated 12-well transwell filters (CLS3460; Sigma-Aldrich) or cover glasses (12-545-81; Thermo Fischer Scientific). For β-gal staining, cells were fixed for 5min at RT in a β-gal fixation solution (5mM EGTA, 2.5mM MgCl2, 0.2% Glutaraldehyde, 1.3% PFA in PBS) and washed for 5 min at RT with β-gal washing buffer (2mM MgCl2, 0.02% NP-40 in PBS). A β-gal staining was performed at 37°C for 3h in 0.02% X-Gal, 5mM K3Fe(CN)6, 5mM K4Fe(CN)6, 2mM MgCl2 in PBS. After staining, astrocytes were fixed again with 4% PFA for 10 min at RT, pH 7.4, and permeabilized with 0.5% Triton X-100 in PBS for 5min. For astrocytes not undergoing β-gal staining, the cells were fixed for 10 min at RT with 4% PFA in PBS, pH 7.4, and permeabilized with 0.5% Triton X-100 in PBS for 5 min. The cells were blocked with 0.5% BSA for 30min and incubated with rat polyclonal antibodies anti-GFAP (1:80; 13-0300; Thermo Fischer Scientific), sheep polyclonal antibodies anti-ITβ5 (1:40, R&D Systems), rabbit polyclonal antibodies anti-HIFα (1:150; NB100-134; Novus Biologicals), and goat polyclonal antibodies anti-SOX-9 (1:40; AF3075; R&D Systems) overnight at RT. Cells were washed four times with PBS and incubated with anti-rabbit Alexa Fluor 594, and anti-rat Alexa Fluor 488 secondary antibodies (1:300; Thermo Fischer Scientific) or anti-Goat Alexa Fluor 594, anti-rabbit Alexa fluor 488 secondary antibodies (1:300; Jackson ImmunoResearch) in PBS for 1h at RT. Astrocyte nuclei were stained with DAPI and mounted with Fluoromount-G mounting medium (SouthernBiotech). Human tissue was obtained after informed consent from patients undergoing lesion resection, under protocol #10-295-A, approved by the University of Chicago Institutional Review Board. Human tissue, CCM lesions, and lesion-free brain tissues were snap-frozen and sectioning using a cryostat (Leica). Specimens were air dry for 15 min and fixed in 4% PFA at room temperature for 15 min and washed three times in PBS. The specimens were blocked and permeabilized using permeabilization buffer for 2 h and incubated with rabbit polyclonal antibody anti-eNOS (1:200; PA1-037; Thermo Fisher Scientific) and goat polyclonal antibody anti–collagen IV (1:100, AB769; Millipore) in PBS at room temperature overnight. Preparations were washed four times in PBS and incubated at room temperature for 1 h with suitable secondary anti-rabbit Alexa Fluor 594 and anti-goat Alexa Fluor 488 antibodies (1:300; Thermo Fisher Scientific) in PBS. Cell nuclei were stained with DAPI (SouthernBiotech).

### Immunohistochemistry

Brains from *Pdcd10^ECKO^* and littermate control *Pdcd10^fl/fl^* mice at postnatal day 10 were isolated and fixed in 4% PFA at 4°C overnight. After cryoprotection in 30% sucrose dissolved in PBS, brains were embedded and frozen in O.C.T compound (23-730-571; Fischer Scientific). Cerebellar tissues were cut into 12-μm coronal sections onto Superfrost Plus slides (12-550-15; VWE International). Sections were blocked and permeabilized in a permeabilization solution (0.5% Triton X-100, 5% goat serum, 0.5% BSA, in PBS) for 2h and incubated in rabbit polyclonal antibodies against eNOS (1:200; PA1-037; Thermo Scientific), rabbit polyclonal antibodies against GFAP (1:250; GA524; Agilent Dako), mouse monoclonal antibody against MBP (1:500; SMI99; Biolegend), rat polyclonal antibodies against CD31 (1:80; 553370; BD Biosciences), rat monoclonal antibody against CD34 (1:100; 119302; Biolegend) in PBS at room temperature overnight. Preparations were washed one time in brain-Pblec buffer (PBS, 1mM CaCl2, 1mM MgCl2, 0.1 mM MnCl2, and 0.1% Triton X-100) and incubated with isolectin B4 FITC conjugated (1:80, L2895; Sigma-Aldrich) in brain-Pblec buffer at 4C overnight. Tissue sections were washed four times in PBS and incubated with suitable Alexa Fluor coupled secondary antibodies (1:300, Thermo Fischer Scientific) in PBS for 1h at RT. Cell nuclei were stained with DAPI and mounted with Fluoromount-G mounting medium (SouthernBiotech). For β-gal staining, brains were fixed for 5min at RT in PFA 2% and washed with PBS. After cryoprotection in 15% sucrose dissolved in PBS, brains were embedded and frozen in O.C.T compound. A β-gal staining was performed at 37°C for 6h in 0.02% X-Gal, 5mM K3Fe(CN)6, 5mM K4Fe(CN)6, 2mM MgCl2 in PBS and tissue sections were washed with cold PBS and fixed in 4% PFA at RT for 30 min. Immunohistochemistry was performed after the β-gal staining. Except for immunohistochemistry for SOX-9 in which a β-gal staining was performed at 37°C for 3h followed by fixation with PFA 4% for 30 min at RT and incubation with unmasking antigen solution (927901; Biolegend). Sections were blocked and permeabilized in a permeabilization solution for 2h and incubated in goat polyclonal antibodies against SOX-9 (1:40; AF3075; R&D Systems) and rabbit polyclonal antibodies against GFAP (1:250; GA524; Agilent Dako) overnight at RT. Tissue sections were washed four times in PBS and incubated with suitable Alexa Fluor coupled secondary antibodies (1:300, Thermo Fischer Scientific) in PBS for 1h at RT. The slides were viewed with a high-resolution SP8 confocal microscope (Leica Microsystems), and the images were captured with Leica application suite software (Leica Microsystems).

### In Situ Hybridisation

Brains from *Pdcd10^ECKO^* and littermate control *Pdcd10^fl/fl^* mice at postnatal day 10 were isolated and fixed in 4% PFA at 4°C overnight. Brains were washed in cold RNAse free PBS and after cryoprotection in 30% sucrose dissolved in RNAse free PBS, brains were embedded and frozen in O.C.T compound (23-730-571; Fischer Scientific) and processed for in situ hybridization (ISH) as described previously(92). Immunohistochemistry was performed after the hybridization using rabbit polyclonal antibodies against GFAP (1:250; GA524; Agilent Dako) overnight at RT. Preparations were washed one time in brain-Pblec buffer and incubated with isolectin B4 FITC conjugated (1:80, L2895; Sigma-Aldrich) in Pblec buffer at 4C overnight. Tissue sections were washed four times in PBS and incubated with suitable Alexa Fluor coupled secondary antibodies (1:300, Thermo Fischer Scientific) in PBS for 1h at RT.

### Western blot analysis

BMECs were lysed using a solution containing 1mM sodium orthovanadate, protease inhibitor cocktail (11836170001; Roche), and 100μL of radioimmunoprecipitation assay buffer (Thermo Scientific). A Micro BCA protein assay kit (500-0116; Thermo Scientific) was used to determine the protein concentration and 25μg of cell lysates were heated for 5 minutes at 95°C to denature all proteins. The cell lysates were subjected to a 4% to 20% gradient sodium dodecyl sulfatepolyacrylamide gel electrophoresis (XP04200, Thermo Fischer Scientific) and a wet transfer was used to transfer proteins to nitrocellulose membranes (Amersham). Membranes were blocked for 1h at RT using a blocking solution (TBS 1x, 10% nonfat milk), and polyclonal rabbit antibodies directed against eNOS (1:500; PA1-037; Thermo Scientific), HIF-1α (1:150; NB100-134; Novus Biologicals) or monoclonal rabbit (1:100; 12282; Cell Signaling) and polyclonal goat (1:100; PA1-9032; ThermoFisher) antibodies directed against COX-2 were incubated at 4°C overnight. Several washes were performed and then the membranes were incubated with appropriate IRDye/Alexa Fluor-coupled secondary antibodies (1:10,000, 926-68070; 926-32211; Li-COR) for 1h at RT. A mouse monoclonal antibody against beta-actin (1:5000; A5441; Sigma-Aldrich) was used as a control for protein loading. Membranes were imaged and analyzed using Odyssey CLx Infrared Imaging (Li-COR).

### RNA isolation

Total RNA from cultured astrocytes and BMECs were isolated by TRIzole as specified by the manufacturer’s protocol (Thermo Fischer Scientific). Briefly, 1 ml of TRIzole reagent was added per well. Cell homogenization was completed by pipetting up and down several times throughout the entire surface area where cells were growing. Cell lysates were transferred to Phase Lock Gel 2ml tubes (2302830; VWR). Then, 200μl of chloroform (ICN19400290; Thermo Fischer Scientific) was added to each tube and mixed vigorously for 15 seconds, followed by a 3-minute incubation at room temperature prior to centrifugation at 12,000g for 15 minutes at 4°C. The aqueous phase containing RNA was collected and transferred to a 1.5ml DNAse/RNAse free microfuge tube. To precipitate the RNA, 500μl of isopropanol was added, resuspended, and incubated for 10 minutes at room temperature followed by centrifugation at 12,000g for 10 minutes at 4°C. The supernatant was removed, and the pellet was washed with 1ml of 75% ethanol followed by centrifugation at 7,500g for 5 minutes at 4°C. The supernatant was removed, and the pellet was air-dried at room temperature and dissolved in 11μl-20μl of DNAse/RNAse free water. To determine the concentration and purity, 1μl of each sample was analyzed using UV spectrophotometry at 260 and 280 nm using NanoDrop 1000 Spectrophotometer.

### RT^2^-qPCR analysis

10ng of total RNA was used to produce single-stranded complementary DNA (cDNA) using random primers according to the manufacturer’s protocol (48190011; 18080093; Thermo Fischer Scientific). Briefly, 10ng of RNA was added to a master mix containing 1X First-Strand buffer, 1μl 0.1M DTT, 1mM dNTPs, 1μl RNaseOUT recombinant RNase inhibitor, SuperScript III reverse transcriptase, and 0.5μg random primers. The mixture was placed in a thermal cycler (C1000 Touch, Bio-Rad) at 25°C for 10 minutes, and then incubated at 50°C for 1 hour followed by inactivation of the reaction by heating at 70°C for 15 minutes. 300ng of cDNA was run with the Kapa SybrFast qPCR mix (Kapa Biosystems) using 50μM of primers to distinguish between the relative levels of genes in each condition. The PCR reaction was placed in a thermal cycler (C1000 Touch, Bio-Rad) using an initial step at 95°C for 15 minutes, followed by 40 cycles (30sec at 95°C, 30sec at 55°C, and 30sec at 72°C). Analysis of the data was performed using the 2^-ΔΔCT^ method.

### NO Determination

*Pdcd10^ECKO^* and *Pdcd10^fl/fl^* BMECs were maintained in BMEC media supplemented with 500μM L-arginine and deficient in fetal bovine serum and ascorbic acid for 36 hours. Then, the BMEC media was collected, and a colorimetric total nitric oxide assay was performed as specified by manufacturers protocol (KGE001; R&D Systems). 50μl of undiluted BMEC media samples and nitrate (NO^3^) standards ranging 100μM to 1.565μM were plated on clear polystyrene microplates (DY990; R&D Systems). To measure total nitrites (NO^2^) and nitrates, 25μl of nitrate reductase and 25μl of NADH were added to each well to convert all available nitrates to nitrite. The plate was covered with an adhesive strip and incubated at 37°C for 30 minutes. After incubation, 50μl of Griess reagent I and Griess reagent II were added to all wells and incubated at room temperature for 10 minutes. A two-step diazotization reaction occurs in which NO_2_- reacts with sufanilic acid to produce a diazonium ion, which is then coupled to N-(1-naphthyl) ethylenediamine to form an azo-derivative that absorbs light at 540nm. The optical density (O.D.) of each well was read using a microplate reader (Infinite 200 PRO, Tecan) at 540nm with a wavelength correction of 690nm. Duplicate readings were averaged and normalized to the blank. A standard curve was generated for each experiment by plotting optical density against concentration (μM), and total nitrate/nitrite concentrations were determined using the linear trendline.

### Statistics

Statistical analysis for single comparisons was performed using a two-tailed student’s *t*-test and analysis for multiple comparisons was performed using ANOVA followed by Tukey’s post hoc test. The threshold for statistical significance was set at a *p*-value of 0.05.

### Study approval

All animal experiments were performed in compliance with animal procedure protocols approved by the University of California, San Diego Institutional Animal Care and Use Committee.

## Author contributions

M.A.L.R., S.I.S., P.H., C.C.L., performed and analyzed histology and gene expression experiments in culture and conditional knockout mice; S.I.S., P.H., C.C.L., Substantial contribution to manuscript preparation, writing, analysis of the data and generation of all figures; A.P., R.V., E.E., S.M., Performed histology and treatment of conditional knockout mice; R.G., T.M., R.L., N.H., I.A.A., Performed, analyzed, and interpreted the microCT studies and manuscript editing. S.M., O.P., G.G.H., R.D., Data discussion and manuscript editing; B.G., H.S., F.L., M.H.G., Substantial contribution to manuscript preparation, writing, and interpretation of data; M.AL.R. Designed the experiments, prepared the figures, and with M.H.G. performed overall study design to address the conceptual ideas, analysis and interpretation of the data, and writing of the manuscript.

## Acknowledgments

The authors thank Angioma Alliance for providing human CCM specimens; Chelsea Hyun Ju Choi, Alyssa Castillo, Daniel Han and Maryum Haidari for technical assistance; Jennifer Santini and Marcy Erb for microscopy technical assistance; and the UCSD School of Medicine Microscopy Core.

## Supplementary Materials

**Fig. S1.**
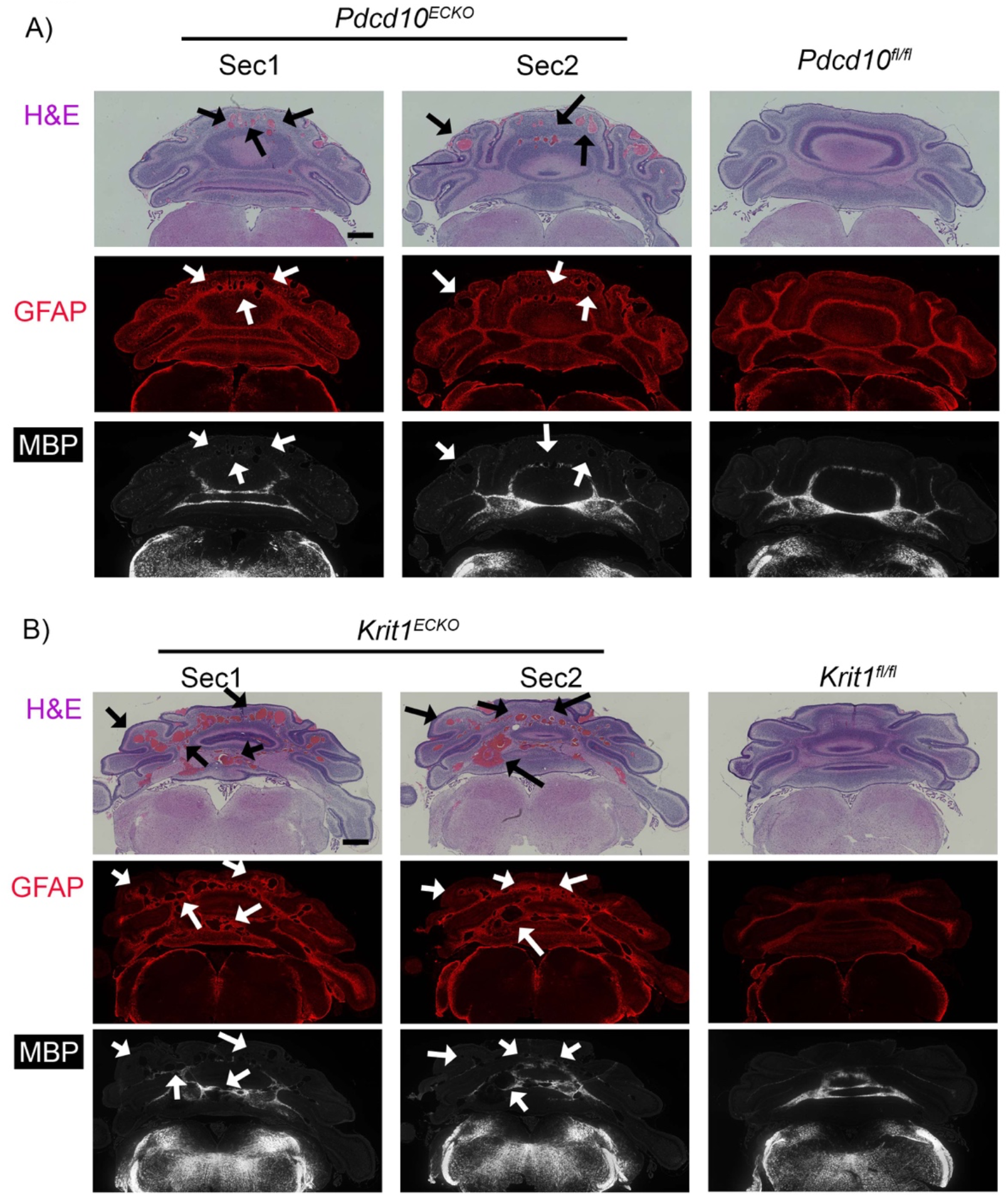
CCM lesions spatially developed on fibrous astrocytes. (**A**)Histological analysis of cerebellar sections from P9 *Pdcd10^ECKO^* and littermate control *Pdcd10^fl/fl^* mice. Low magnification of CCM lesions detected in sections stained by hematoxylin and eosin. CCM lesions spatially developed on fibrous astrocytes areas positive to GFAP immunostaining (red) and white matter positive to MBP (white). Arrows indicate CCM lesions. (**B**) Histological analysis of cerebellar sections from P8 *Krit1^ECKO^* and littermate control *Krit1^fl/fl^* mice. Low magnification of CCM lesions detected in sections stained by hematoxylin and eosin. CCM lesions spatially developed on fibrous astrocytes areas positive to GFAP immunostaining (red) and white matter positive to MBP (white). Arrows indicate CCM lesions. Scale bar:(**A** and **B**) 500 μm.

**Fig. S2.**
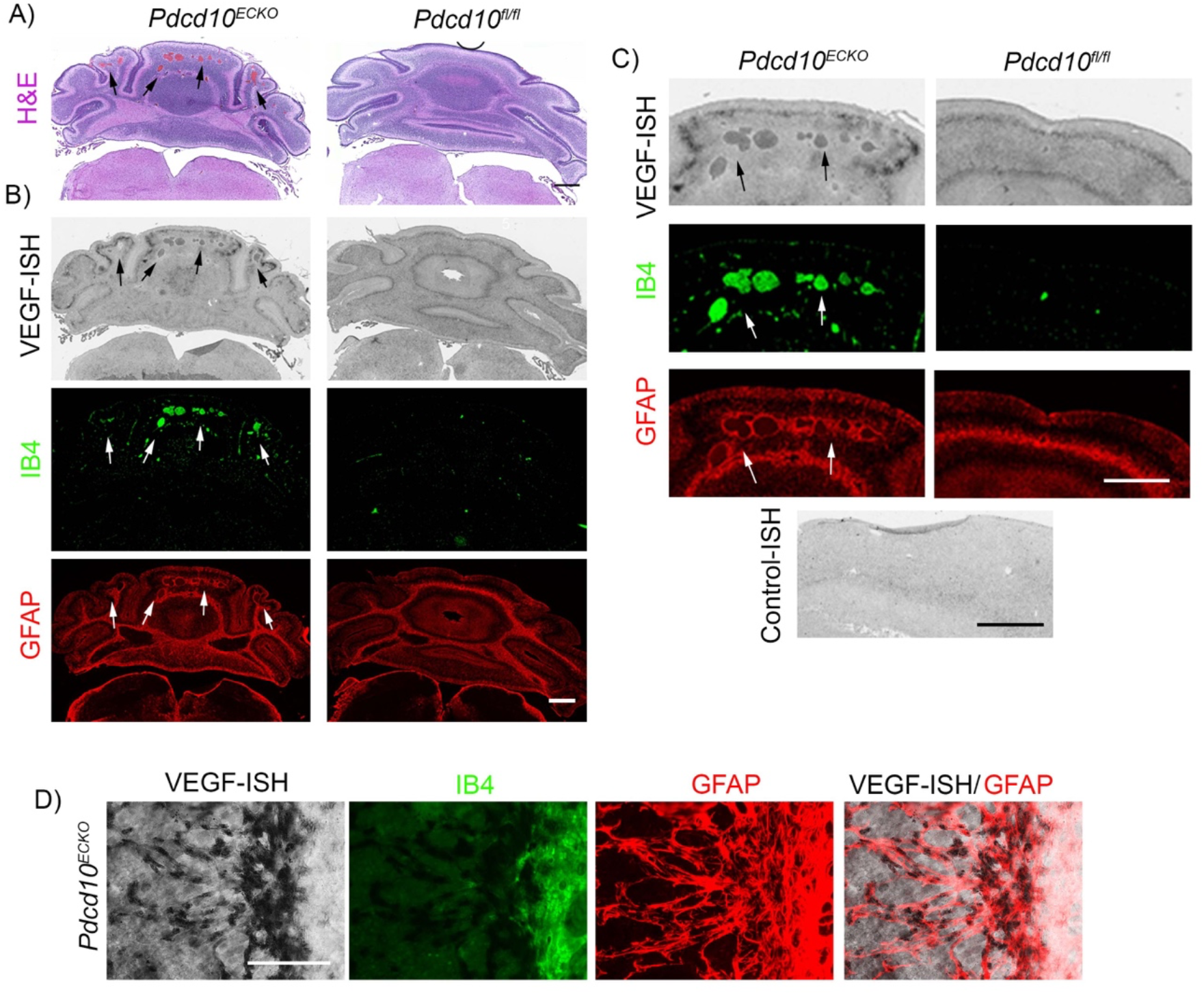
VEGF increases during cerebral cavernous malformations development. (**A**)Histological analysis of cerebellar sections from P10 *Pdcd10^ECKO^* and littermate control *Pdcd10^fl/fl^* mice. Low magnification of CCM lesions detected (Arrows) in sections stained by hematoxylin and eosin. (**B**) ISH for VEGF (black) combined with immunohistochemistry to identify GFAP-positive astrocytes (red), endothelial marker isolectin B4 (IB4; green) in a serial section from *A*. (**C**) high magnification of ***B*** and control probe for ISH. (**D**) ISH for VEGF (black) combined with immunohistochemistry to identify GFAP-positive astrocytes (red), endothelial marker isolectin B4 (IB4; green) in P10 *Pdcd10^ECKO^* retinas (n=2 or 3). Scale bars: (**A**, **B** and **C**) 500 μm, (**D**) 100 μm.

**Fig. S3.**
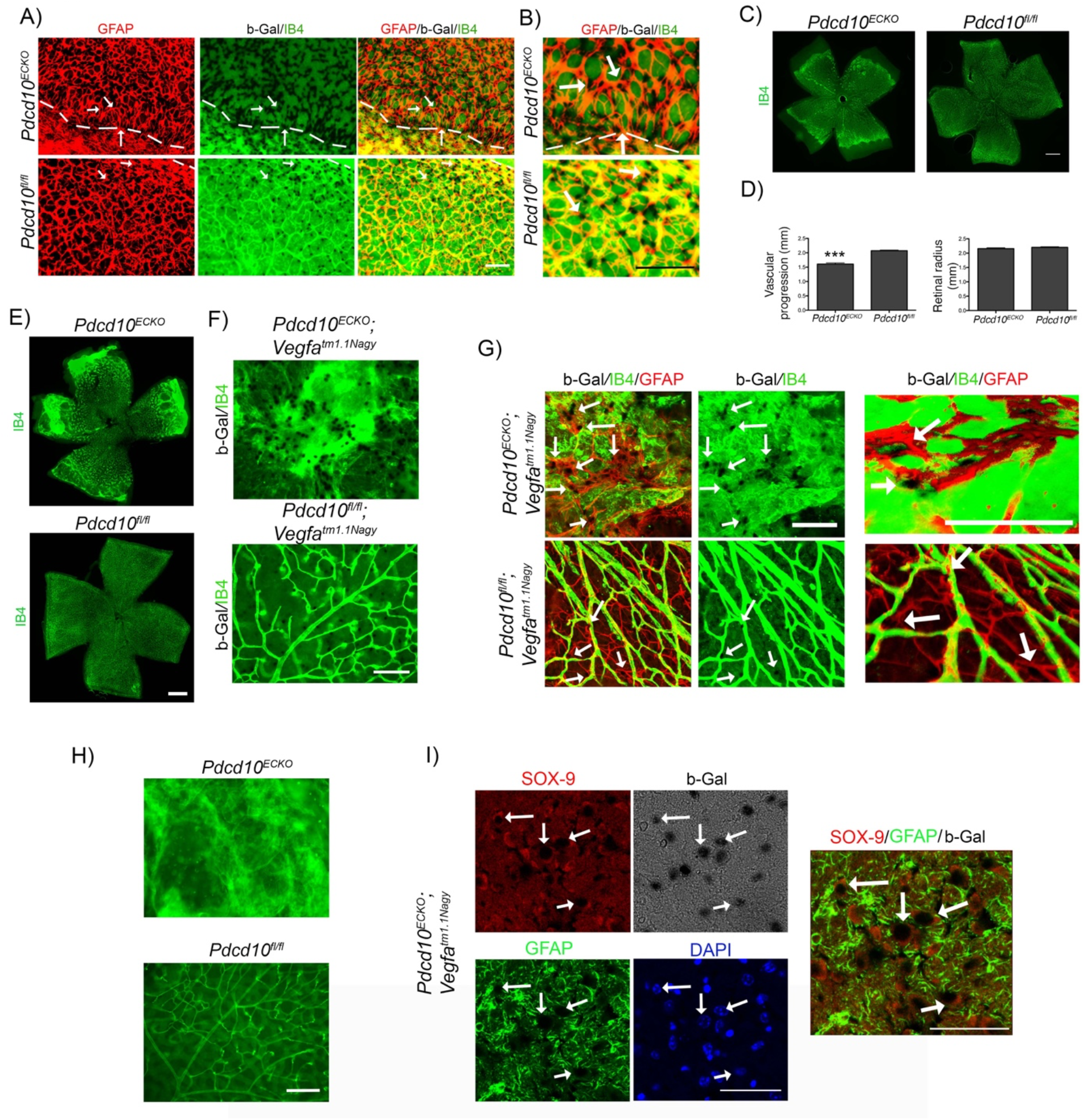
VEGF increases in retinas during cerebral cavernous malformations. (**A**) Wholemount retinal vasculature at the angiogenic growth front stained for GFAP (red), Isolectin B4 (green), and β-gal/VEGF expression detected by X-gal staining (black) in P9 *Pdcd10^ECKO^;Vegfa^tm1.1Nagy^* mice and *Pdcd10^fl/fl^;Vegfa^tm1.1Nagy^* littermate control. Dotted line indicated angiogenic front. (**B**) Magnified whole-mount retinal vasculature in *A*. Arrows indicate β-gal/VEGF in GFAP-positive astrocytes. (**C**) Isolectin B4-stained *Pdcd10^ECKO^* and control *Pdcd10^fl/fl^* P9 retinas and in (**D**) the quantification of vascular parameters of retinas (SEM, *n*= 14 or 17 mice in each group). (**E**) Whole-mount retinal vasculature by isolectin B4-stained *Pdcd10^ECKO^* and control *Pdcd10^fl/fl^* P12 retinas. (**F**) Magnified whole-mount retinal vasculature indicate β-gal/VEGF expression in P12 *Pdcd10^ECKO^;Vegfa^tm1.1Nagy^* mice and *Pdcd10^fl/fl^;Vegfa^tm1.1Nagy^* littermate control. (**G**) Correspond to β-gal control in P12 *Pdcd10^ECKO^* mice and *Pdcd10^00^* littermate control. (**H**) Maximum-intensity projection of whole-mount P12 retinal vasculature from *F* stained for GFAP (red), Isolectin B4 (green), and β-gal/VEGF (black). Arrows indicate β-gal/VEGF in GFAP-positive astrocytes. Surface reconstruction of β-gal/VEGF in GFAP-positive astrocytes (n=3 or 4). (**E**) Confocal microscopy of cerebrum cortex from *Pdcd10^ECKO^;Vegfa^tm1.1Nagy^* stained for β-gal/VEGF expression, SOX-9-positive astrocytes (red), GFAP-positive astrocytes (green), and DAPI for nuclear DNA (blue). Data are mean±SEM. ***, P<0.001; determine by Student’s *t* test. Scale bars: (**A**, **B** and **H**) 100 μm; (**F** and **G**) 200 μm; (**C** and **E**) 500 μm.

**Fig. S4.**
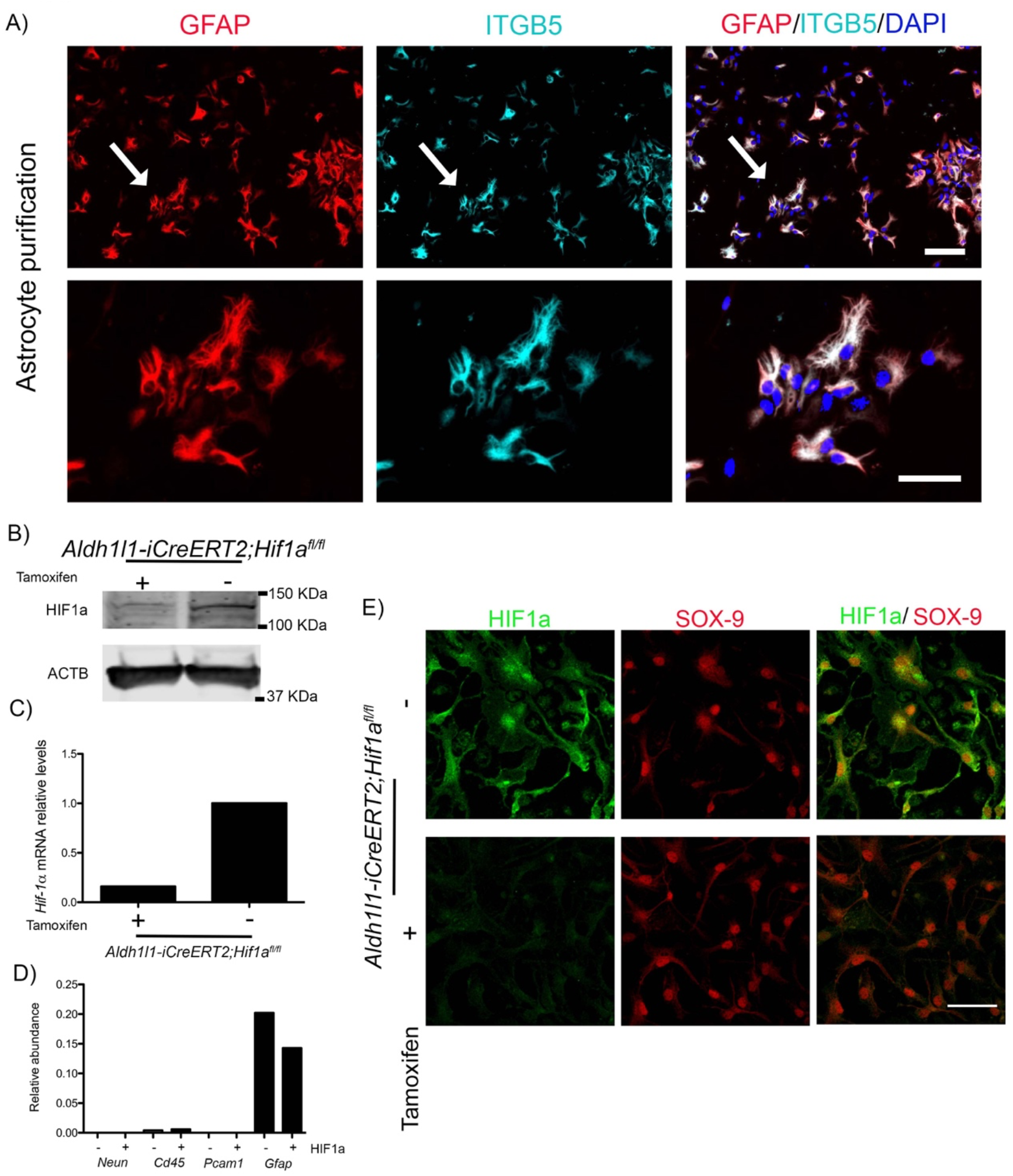
Mouse primary astrocyte culture characterization and purity. Astrocytes isolated from cortex of postnatal day 5-7 mice. (**A**) Purity of the isolation was confirm by the presence of GFAP and ITGB5 astrocyte markers visualized by immunofluorescence (n=5). (**B**) the HIF-1α antibody specificity by Western blot analysis of primary astrocyte cultures was determined using *Aldh1l1-iCreERT2;Hif-1 α^fl/fl^* astrocytes in the presence and absence of tamoxifen (n=2). (**C**) Inactivation of HIF-1α was further validated by RT-qPCR that showed 80% reduction in HIF-1α mRNA levels in *tamoxifen-treated-Aldh1l1-iCreERT2; Hif-1α^fl/fl^*cells (n=2). (**D**) Gene expression of neuronal marker *Neun;* leucocyte marker *Cd45;* endothelial cell marker *Pcam1;* astrocyte markers *Gfap* from *C* (n=2). (**E**) HIF-1α antibody specificity by immunocytochemistry of astrocytes was performed by increasing HIF-1α expression by adding 100 μM of CoCl2 for 24 h to the culture medium. SOX-9 immunostaining was used as a nuclear marker for astrocytes. Diffuse and nuclear accumulation of HIF-1α immunostaining in non-tamoxifen *treated-Aldh1l1-iCreERT2; Hif-1α^fl/fl^* cells that is significantly reduced by tamoxifen treatment astrocytes (n=2). Scale bars: (**A** and **E**) 100 μm.

**Fig. S5.**
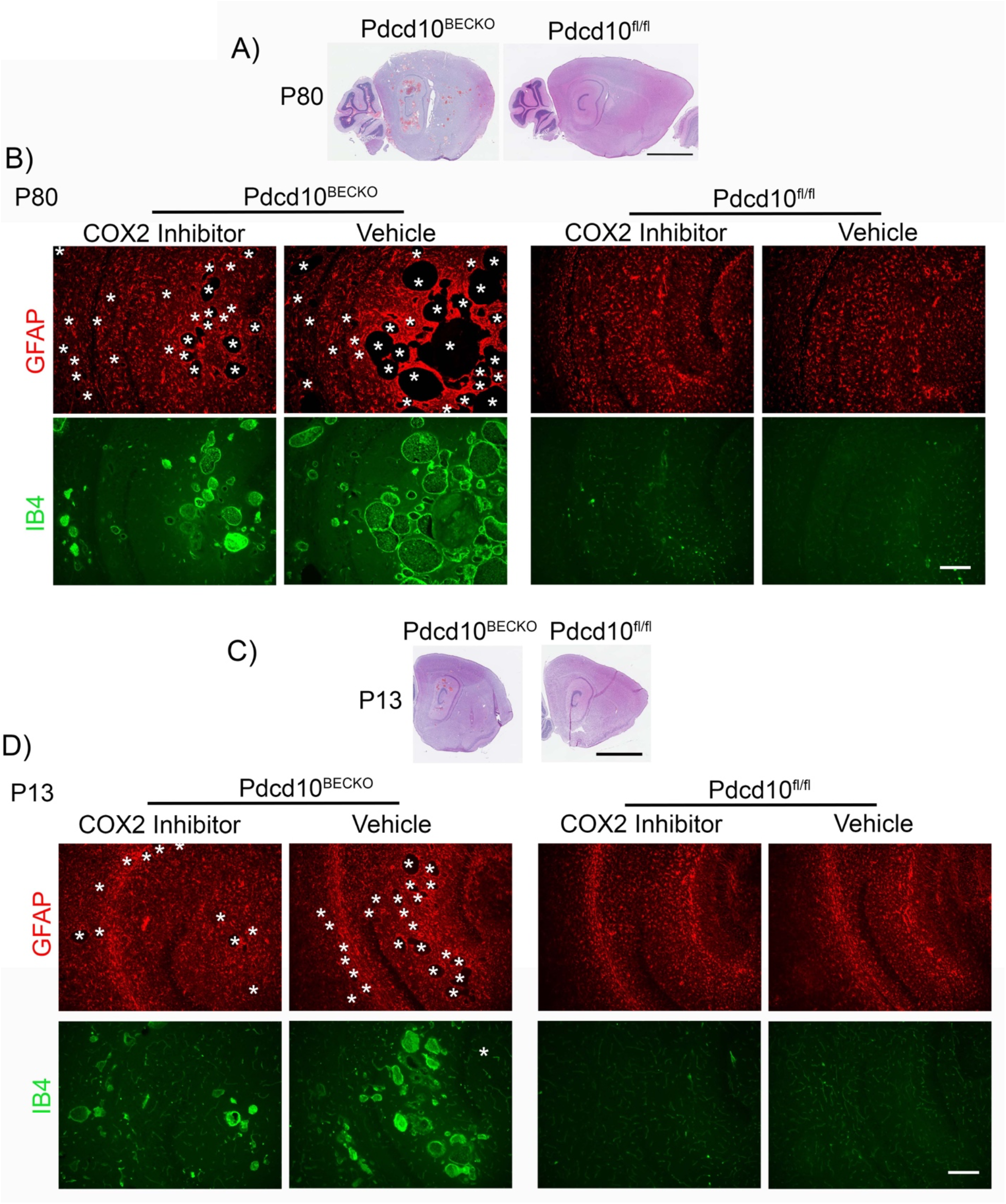
COX-2 inhibition prevent CCM lesions in acute and chronic CCM mouse models. (**A**)Histological analysis of brain sections from P80 *Pdcd10B^ECKO^* and littermate control *Pdcd10^fl/fl^* mice. Low magnification of CCM lesions detected throughout the brain sections stained by hematoxylin and eosin in P80 *Pdcd10B^ECKO^* mice. More extensive and complex lesions were prevalent in the hippocampal area (n=3). (**B**) Oral gavage administration of 40 mg/Kg celecoxib or vehicle for fifteen consecutive days P55 to P70. GFAP astrocyte marker (red) staining and IB4 endothelial marker (green) of mouse hippocampal region at P80. CCM lesions’ high propensity to develop surrounded by GFAP+ astrocytes in the hippocampal region in vehicle-treated *Pdcd10B^ECKO^* mice. Significant decrease in CCM lesions’ density and GFAP-immunoreactivity in celecoxib-treated *Pdcd10B^ECKO^* mice. Arrows indicate CCM lesions (n=3). (**C**)Histological analysis of brain sections from P13 *Pdcd10^BECKO^* and littermate control *Pdcd10^fl/fl^* mice. Low magnification of CCM lesions detected throughout the brain sections stained by hematoxylin and eosin in P13 *Pdcd10B^ECKO^* mice. Extensive lesions were prevalent in the hippocampal area (n=3). (**D**) Intragastric administration of 40 mg/Kg celecoxib or vehicle for four consecutive days P6 to P9. GFAP astrocyte marker (red) staining and IB4 endothelial marker (green) of mouse hippocampal region at P13. CCM lesions’ high propensity to develop surrounded by GFAP+ astrocytes in the hippocampal region in vehicle-treated *Pdcd10B^ECKO^* mice. Significant decrease in CCM lesions’ density in celecoxib-treated *Pdcd10B^ECKO^* mice (n=3). Asterisks, vascular lumen of CCM lesions. Scale bars: (**B** and **D**) 200 μm.

**Fig. S6.**
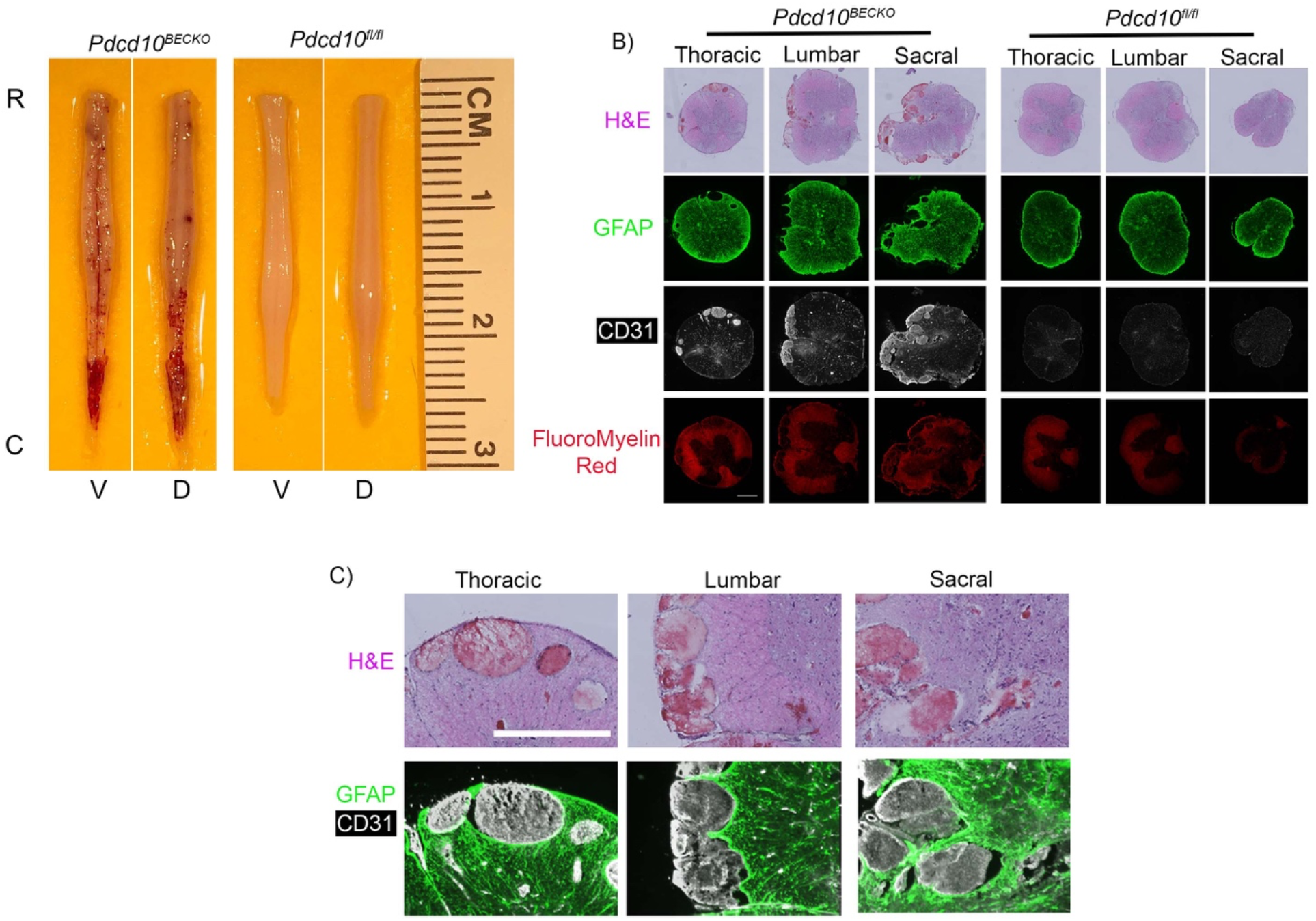
CCM lesions in the spinal cord of chronic CCM mouse model. (**A**) Prominent lesions are present along the spinal cord of P80 *Pdcd10^BECKO^* mice. R=rostral, C=caudal, V=ventral, D=dorsal (n=14). (**B**) Histological analysis of serial sections of spinal cords from P80 *Pdcd10^BECKO^* and littermate control *Pdcd10^fl/fl^* mice. Spinal cord sections stained by hematoxylin and eosin, GFAP astrocyte marker (green), CD31 endothelial marker (white), and myelin staining (red) of a mouse at P80 (n=2). (**C**) High magnification of CCM lesions, in ***B***, present in the thoracic, lumbar, and sacral region of a spinal cord, shown a high propensity to develop surrounded by GFAP+ astrocytes (n=2). Scale bars: (**B** and **C**) 500 μm.

**Fig. S7.**
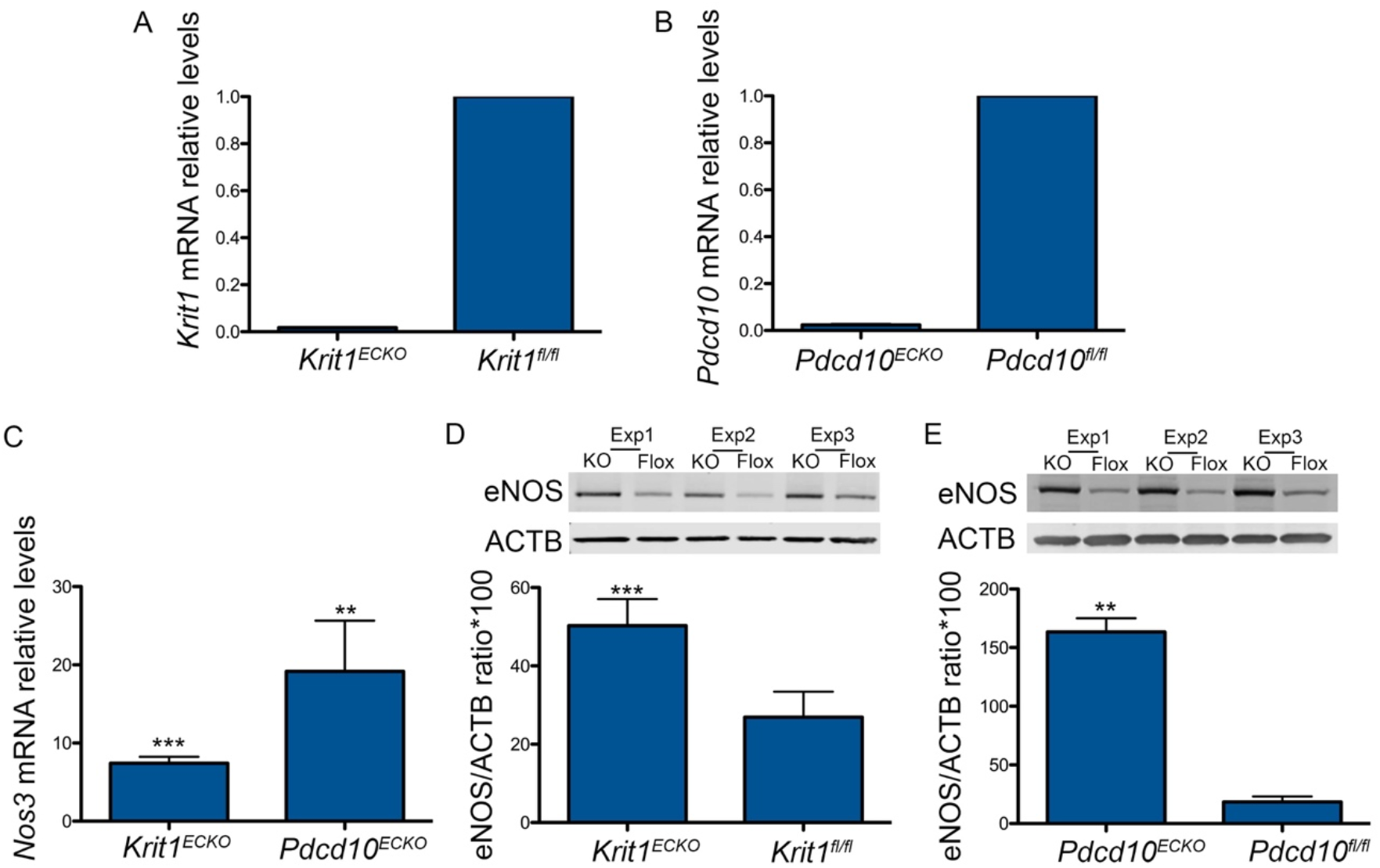
Loss of brain endothelial *Krit1* or *Pdcd10* increases the expression of eNOS. (**A**) Analysis of *Krit1* mRNA levels by RT-qPCR in *Krit1^ECKO^* BMEC and *Krit1^fl/fl^*BMEC control (SEM, *n*=3). (**B**) Analysis of *Pdcd10* mRNA levels by RT-qPCR in *Pdcd10^ECKO^* BMEC and *Pdcd10^fl/fl^* BMEC control (SEM, *n*=3). (**C**) Analysis of *Nos3* mRNA levels by RT-qPCR in *Krít1^ECKO^* BMEC and *Pdcd10^ECKO^* BMEC, as compared to *Krit1^fl/fl^* BMEC or *Pdcd10^fl/fl^* BMEC control, respectively (SEM, n=3). (**D**) Quantification of eNOS protein in *Krit1^ECKO^* BMEC compared with *Krit1^fl/fl^* BMEC control (SEM, n=3). (**E**) Quantification of eNOS protein in *Pdcd10^ECKO^* BMEC compared with *Pdcd10^fl/fl^* BMEC control (SEM, n=3). Data are mean±SEM. **, P<0.01, ***, P<0.001; determine by Student’s *t* test.

**Fig. S8.**
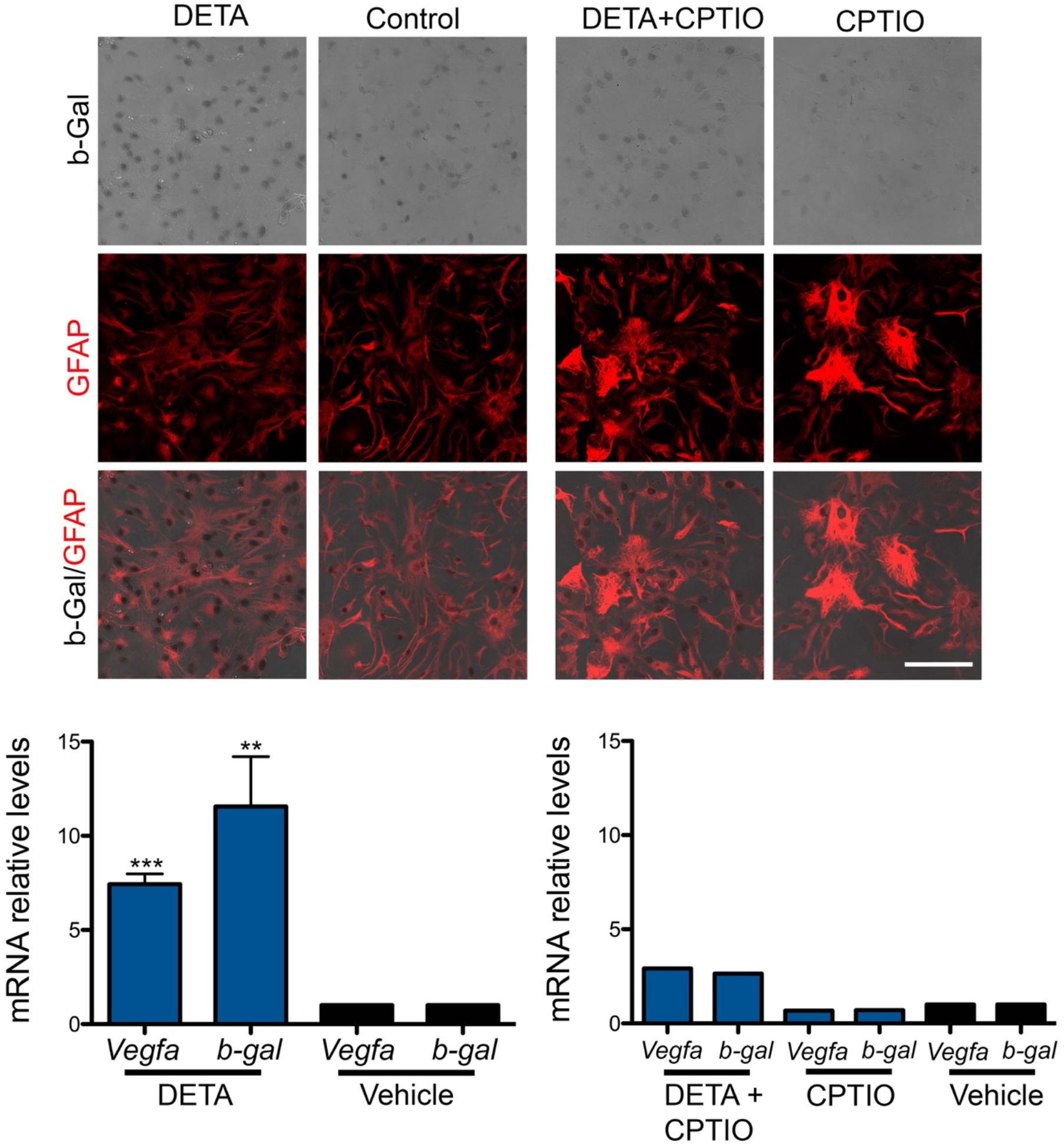
Primary cultures of astrocytes respond to elevation of NO. (**A**) Increase in β-gal/VEGF expression, as shown by X-gal staining (black), in primary cultured astrocytes (GFAP-positive cells, red) treated with 0.5mM DetaNONOate (NO donor) for 24h when compared with vehicle-treated astrocytes. Astrocytes pre-treated with 15μM CPTIO, a NO scavenger, prevented DetaNONOate-induced increase in β-gal/VEGF expression in astrocytes. (**B**) DETANONOate in astrocyte culture media induced an ~7 fold increase in astrocyte *Vegfa* mRNA and an ~11.50 fold increase in astrocyte *β-gal* mRNA levels, an effect that was prevented in astrocytes pre-treated with CPTIO. Data are mean±SEM. **, P<0.01, ***, P<0.001; determine by Student’s *t* test.

